# Pioneer factor GAF cooperates with PBAP and NURF to regulate transcription

**DOI:** 10.1101/2020.05.10.087262

**Authors:** Julius Judd, Fabiana M. Duarte, John T. Lis

**Affiliations:** Department of Molecular Biology and Genetics, Cornell University, Ithaca, New York 14835, USA; Department of Stem Cell and Regenerative Biology, Harvard University, Cambridge, MA 02138, USA

## Abstract

Transcriptionally silent genes must be activated throughout development. This requires nucleosomes be removed from promoters and enhancers to allow transcription factor binding (TFs) and recruitment of coactivators and RNA Polymerase II (Pol II). Specialized pioneer TFs bind nucleosome-wrapped DNA to perform this chromatin opening by mechanisms that remain incompletely understood^1–3^. Here, we show that GAGA-factor (GAF), a *Drosophila* pioneer factor^4^, interacts with both SWI/SNF and ISWI family chromatin remodelers to allow recruitment of Pol II and entry to a promoter-proximal paused state, and also to promote Pol II’s transition to productive elongation. We found that GAF functions with PBAP (SWI/SNF) to open chromatin and allow Pol II to be recruited. Importantly this activity is not dependent on NURF as previously proposed^5–7^; however, GAF also functions with NURF downstream of this process to ensure efficient Pol II pause release and transition to productive elongation apparently through its role in precisely positioning the +1 nucleosome. These results demonstrate how a single sequence-specific pioneer TF can synergize with remodelers to activate sets of genes. Furthermore, this behavior of remodelers is consistent with findings in yeast^8–10^ and mice^11–13^, and likely represents general, conserved mechanisms found throughout Eukarya.

---

Pioneer transcription factors are a class of transcription factors that can bind and open condensed chromatin. They control cell-fate decisions in development by opening chromatin at previously inactive lineage-specific promoters and enhancers via sequence-specific binding^1–3^. These factors possess the unique ability to bind nucleosome-wrapped DNA, but the question of how they evict nucleosomes and initiate transcription remains open.

From yeast to mammals, there is growing evidence that pioneer factors cooperate with multiple ATP-dependent nucleosome remodeling complexes to establish transcription-permissive chromatin architecture^8^. In yeast, the pioneer factor Abf1 synergizes with the RSC complex (SWI/SNF family) to maintain the nucleosome-free region (NFR) of Abf1-bound promoters, while ISW1a and ISW2 are required to properly position the +1 nucleosome and phase downstream nucleosomes^9^. In mouse embryonic stem cells, the pioneer factors OCT4 and NANOG are codependent on BAF complex (SWI/SNF family) subunit BRG1 to bind and open chromatin at target sites^11,13^. Recent structural studies have illuminated how SWI/SNF family remodelers bidirectionally evict nucleosomes from promoter NFRs in yeast^10^ and mammals^12^.

GAGA-factor (GAF) is a *Drosophila* transcription factor encoded by the *Trithorax-like* (*trl*) gene^14^ that preferentially binds GAGAG repeats, but is capable of binding a single GAG trinucleotide^15^. We have previously demonstrated that, in *Drosophila* cell cultures, GAF is essential for establishing paused Pol II on GAF-bound promoters, and that the NFRs of these promoters fill with nucleosomes upon GAF depletion^16^. Without this activity, the response of a subset of heat shock genes is impaired^17^. In early fly embryos, regions with chromatin signatures similar to those at binding sites of the embryonic pioneer factor Zelda—but lacking Zelda binding—are enriched for GAF binding, suggesting that GAF may also be an additional early embryonic pioneer factor^4^. GAF interacts physically with the NURF complex (<u>Nu</u>cleosome <u>R</u>emodeling <u>F</u>actor) and both GAF and NURF are required to remodel nucleosomes on the *hsp70* promoter *in vitro*^5,18^. We have previously speculated that GAF recruits NURF to target promoters, which clears them of nucleosomes and allows Pol II initiation and subsequent pausing to proceed^7^. However, early studies speculated that GAF can also interact with Brahma (Brm) complexes (SWI/SNF family; BAP/PBAP)^6^, and recent evidence indicates that GAF physically interacts with PBAP (Polybromo associated Brm) but not BAP^19,20^ in addition to NURF.

To test which of these remodelers is responsible for GAF’s ability to establish transcription-permissive chromatin architecture at target genes, we depleted GAF, NURF301, and BAP170 (unique subunits of the NURF and PBAP complexes that are essential for complex functionality)—as well as NURF301 and BAP170 simultaneously—in S2 cells using RNAi (Fig. 1a). After confirming knockdown efficiency (Extended Data Fig. 1), we used a combination of PRO-seq, ATAC-seq, and 3’RNA-seq to monitor changes in nascent transcription, chromatin state, and mRNA output.

**Figure 1:**
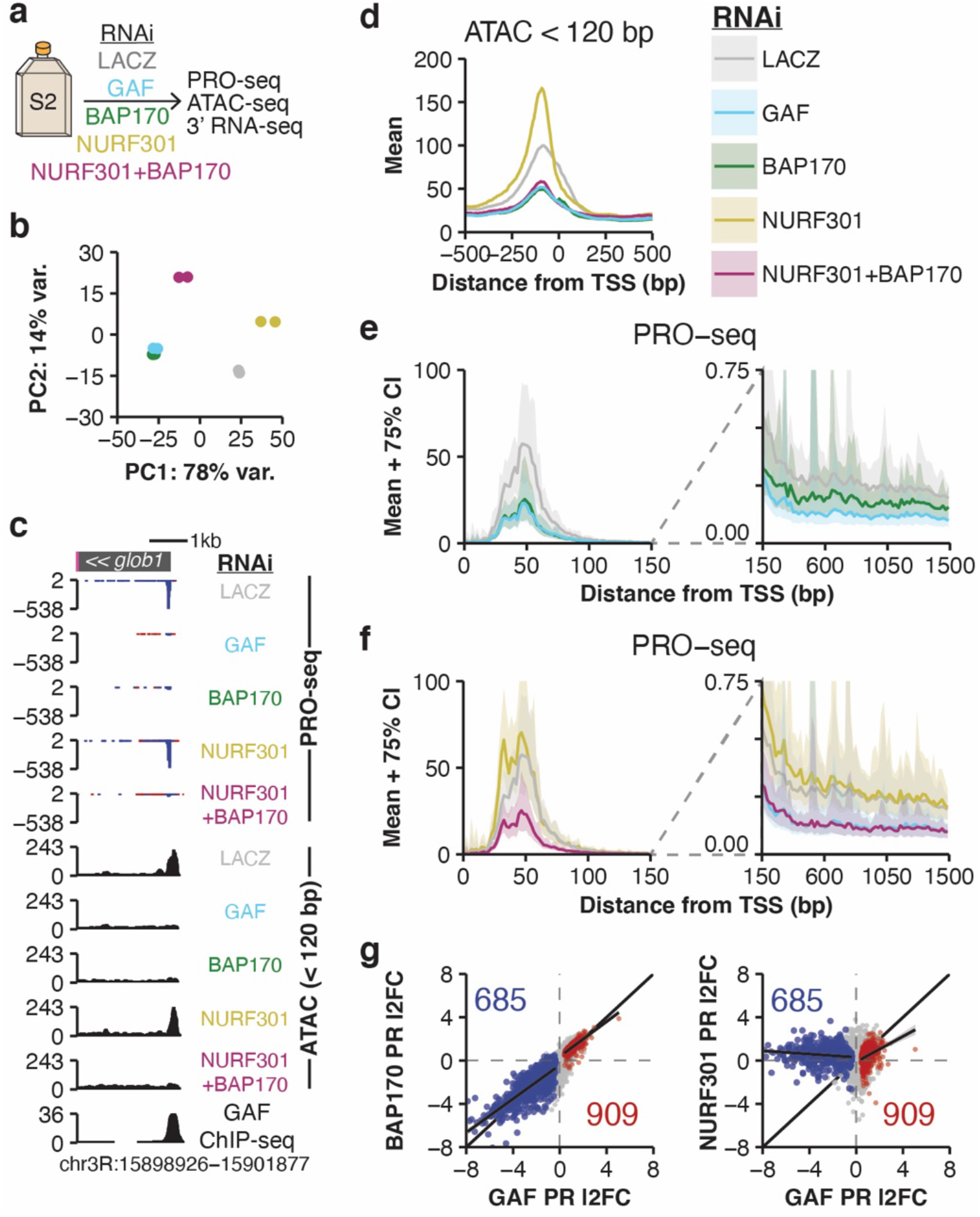
GAGA-factor opens chromatin via the PBAP complex. **(a)** Experimental design. **(b)** Principal component analysis of spike-in normalized PRO-seq signal in the pause region (TSS – 50 to +100). **(c)** Browser shot of *glob1-RB*. **(d)** ATAC-seq (< 120 bp) signal in 1 bp bins at promoters with GAF-RNAi downregulated pausing (n=685; see Extended Data Figure 5a; DESeq2 FDR < 0.01). Signal is the mean of 1,000 sub-samplings of 10% of regions. **(e)** PRO-seq signal for the LACZ, GAF, and BAP170 RNAi treatments. The pause region (left) is in 2 bp bins, and the gene body (right) is in 20 bp bins. Data is shown as mean (line) ± 75% confidence interval (shaded) from 1,000 sub-samplings of 10% of regions. Gene set as in **(d)**. **(f)** As in **(e)**, but for LACZ, NURF301, and NURF301+BAP170 RNAi treatments. GAF-RNAi is also shown in the gene body region for comparison (blue line), though it is partially obscured by the NURF+PBAP line (purple) due to similarity of the trace. **(g)** Pause region (TSS −50 to +100) PRO-seq log2 fold change (l2FC) vs. the LACZ-RNAi control; GAF-RNAi compared to BAP170-RNAi (left) or NURF301-RNAi (right). Red/blue points are significantly changed by GAF-RNAi (DESeq2 FDR < 0.01). Also shown: a GLM and 95% confidence interval for up- and down-regulated promoters.

## GAF synergizes with PBAP to open chromatin

We used a spike-in normalization strategy for PRO-seq and 3’RNA-seq (see methods) to ensure the detection of widespread transcriptional changes that can be hidden by centralizing normalization strategies such as RPKM^21^. A principal component analysis of all genome-wide data sets revealed that GAF-knockdown predominantly clusters with PBAP-knockdown (Fig. 1b, Extended Data Fig. 2). After confirming data quality (Extended Data Fig. 2–4), we defined a set of promoters that have downregulated pause region PRO-seq signal upon GAF knockdown (Extended Data Fig. 5a). Notably, the number of genes with GAF-dependent pausing was far greater than previously reported because our spike-in normalization scheme allowed us to examine the genome-wide effects of GAF depletion with unprecedented sensitivity (n=685 in this study, n=140 reported previously^16^). ATAC-seq hypersensitivity signal (fragments < 120 bp, see methods) revealed that these promoters are substantially less accessible upon GAF, PBAP, or NURF+PBAP knockdown (Fig. 1c–d, Extended Data Fig. 6), and PRO-seq shows that pausing is severely reduced upon GAF, PBAP, or NURF+PBAP knockdown, but not after NURF knockdown (Fig. 1c, e, f, Extended Data Fig. 6). These results clearly demonstrate that GAF coordinates with PBAP–not NURF as previously proposed–to regulate Pol II recruitment by evicting nucleosomes from the NFRs of target promoters. To our knowledge, this is the first report of a pioneer factor synergizing specifically with PBAP (or PBAF, the homologous mammalian complex) to maintain accessible target promoters in metazoans.

In contrast to PBAP, NURF knockdown increases PRO-seq signal in the pause region and in the gene body region compared to the LACZ-RNAi control, particularly in the early pause region closer to the TSS (Fig. 1f, left panel). We then compared the changes in pause region PRO-seq signal upon GAF knockdown to that observed after PBAP and NURF knockdown on a gene-by-gene basis. This revealed a near-perfect one-to-one correlation between GAF and PBAP knockdowns, but minor anticorrelation between GAF and NURF knockdowns (Fig. 1g, compare left panel to right panel, and Extended Data Fig. 7d for the NURF+PBAP knockdown). When we examined promoters with PBAP-dependent pausing (n=806; Extended Data Fig 5b), we observed similar trends to those seen at GAF-dependent promoters: decreased pausing and promoter accessibility after GAF, PBAP, and NURF+PBAP knockdown, and increased pausing and narrowed promoter accessibility upon NURF knockdown (compare Extended Data Fig. 7a–c to Fig. 1d-f). Taken together, these data indicate that PBAP and GAF act together to free the promoter of nucleosomes, while NURF acts at a downstream step.

Is GAF’s mechanistic role to bind nucleosome-bound DNA and recruit the PBAP remodeling complex where they act synergistically to remove nucleosomes, or does GAF binding have an intrinsic ability to displace nucleosomes? The striking loss of PRO-seq signal and loss of chromatin openness of promoters (Fig. 1c–d) described when either factor is depleted argues for a highly synergistic model, where GAF alone has little intrinsic chromatin opening activity. To investigate further, we compared ATAC-seq signal between the GAF and PBAP knockdown conditions, which revealed significant low-magnitude changes at only a small number of sites (Extended Data Fig. 8a), indicating that GAF does not possess sufficient intrinsic chromatin opening ability to account for the effects of GAF knockdown on chromatin. In further support of this, 88% of promoters with decreased pausing upon GAF knockdown had decreased pausing upon PBAP knockdown (n=603; Extended Data Fig. 8b).

Small sets of promoters show only GAF- or only PBAP-knockdown effects. We found that GAF-specific promoters (n=82; PBAP knockdown causes no change) had higher levels of GAF ChIP-seq signal and were far less sensitive to the GAF-knockdown than the class of genes dependent on both GAF and PBAP (Extended Data Fig. 8c-d). We speculate that these promoters may be held open by paused Pol II^22^ that is generated by mechanisms independent of PBAP, or the level of PBAP remaining after knockdown was sufficient to be recruited by the high level of GAF bound at these promoters. PBAP-specific promoters are mostly not bound by GAF (Extended Data Fig. 8c), and often contain the binding motif for the transcription factor lola (Extended Data Fig. 8f), which might function like GAF in its collaboration with PBAP.

## M1BP can establish paused Pol II independent of GAF

Not all GAF bound promoters have GAF-dependent pausing, and some of these bound but unaffected promoters are bound by M1BP (<u>M</u>otif <u>1</u> <u>B</u>inding <u>P</u>rotein) and the insulator BEAF-32 (<u>B</u>oundary <u>E</u>lement <u>A</u>ssociated <u>F</u>actor)^16,23^. However, it remains unclear whether M1BP acts redundantly with GAF to open promoters and promote pausing, or if M1BP/BEAF-32 simply insulate promoters from GAF’s activity. To investigate this, we divided GAF-bound promoters into two classes based on whether they have GAF-dependent pausing (n=600) or not (n=1,245), and found that the BEAF-32 and M1BP motifs^23,24^ were overrepresented in GAF-bound promoters with unchanged pausing (Fig. 2a). We then subdivided the class of GAF-bound, GAF-independent pausing genes by whether they were bound by M1BP^23^ (n=152), BEAF-32^25^ (n=152), or both (n=159; Fig. 2b). GAF-binding was weaker and more diffuse in GAF-bound genes with GAF-independent pausing (Class II–IV), while these promoters were directly and strongly bound by either M1BP or BEAF-32 or both (Class II–IV). We know from our previous study that genes bound by M1BP have reduced pause region PRO-seq signal upon M1BP knockdown^17^, and this reduction in pausing correlates with M1BP binding intensity (Fig. 2b). Moreover, all classes of GAF-bound, GAF-independent pause genes had relatively unchanged ATAC-seq hypersensitivity signal in promoters after GAF or BAP knockdown (Fig 2b, left). This demonstrates that M1BP can open chromatin at promoters and create paused Pol II independent of GAF/PBAP, and the weak and diffuse GAF binding at these sites is insufficient to complement depletion of M1BP.

**Figure 2:**
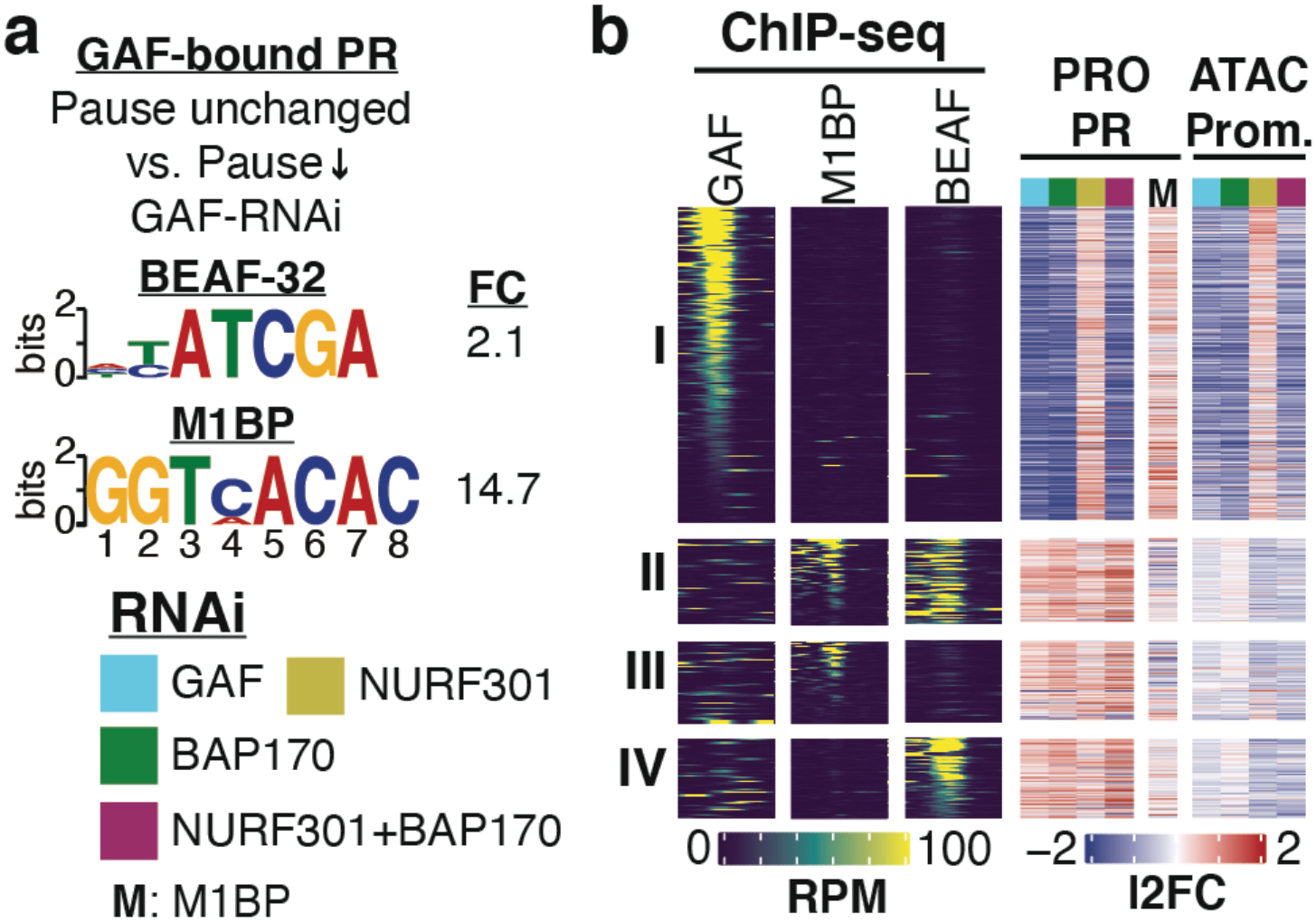
M1BP and BEAF-32 block GAFs activity. **(a)** Motifs enriched in GAF-bound promoters (GAF ChIP-seq peak within −500–TSS) with GAF-independent pausing (n=1,245) over GAF-bound promoters with GAF-dependent pausing (n=600). FC: fold change, DREME E-value < 0.001. **(b)** GAF, M1BP, and BEAF-32 ChIP-seq signal in 10 bp bins in the promoter region (left, TSS ± 500), pause region (TSS −50 to +100) PRO-seq log2 fold change (middle), and promoter (−250 to TSS) ATAC-seq (< 120 bp) log2 fold change (right) at all GAF-bound genes. **(I)** GAF-dependent pausing (n=600); **(II–IV)** GAF-independent pausing (n=1,245); **(II)** M1BP and BEAF-32 bound (n=159); **(III)** M1BP only (n=152); **(IV)** BEAF-32 only (n=152). Sort order: **(I)** GAF ChIP-seq; **(II–III)** M1BP ChIP-seq; **(IV)** BEAF-32 ChIP-seq. GAF-bound GAF-independent genes without M1BP or BEAF-32 ChIP-seq signal are not shown (n=782).

## NURF promotes transition to productive elongation

GAF can physically interact with the remodelers PBAP and NURF^5,19,20^ and appears to function with each remodeler at distinct steps in transcription: GAF and PBAP open chromatin allowing Pol II initiation and entry to the promoter-proximal pause site; while GAF and NURF ensure efficient transition to productive elongation. This role of GAF and PBAP in the first of these two steps is supported strongly by results described above (Fig 1). Evidence that NURF’s role is downstream of PBAP is provided by the observation that the PBAP+NURF double-knockdown primarily mimics PBAP depletion in terms of changes in ATAC-seq and PRO-seq patterns in the pause region (Fig 1). Support for NURFs role in productive elongation comes in part from the fact that the PBAP knockdown only partially recapitulates the decrease in gene body polymerase density seen after GAF depletion (Fig. 1e, right panel). In contrast, the NURF+PBAP double-knockdown mirrors the GAF knockdown (Fig. 1f, right panel). Furthermore, CUT&RUN assays demonstrate co-occupancy of GAF and NURF at promoters genome-wide (Extended Data Fig. 9a), indicating GAF and NURF are likely to act together. These results support the model that GAF coordinates with both remodelers to ensure efficient transcription by first acting with PBAP to open chromatin and allow for the formation of promoter-proximal paused Pol II, and then with NURF to establish chromatin structure at the start of genes which ensures proper transition to productive elongation by Pol II.

How mechanistically can NURF contribute to productive elongation? Knockdown of NURF alone leads to increased highly proximal pausing on a set of promoters (n=831; Extended Data Fig. 5c) and this is coupled with improper +1 nucleosome positioning and phasing of early gene body nucleosomes at these promoters (Fig. 3a). We interpret this decrease in signal at the +1 nucleosome as misphasing, because less consistent positioning would lead to a decrease in aggregated signal at the dyad. While ATAC-seq is not the most precise method of mapping nucleosomes, in light of NURF’s known activity of sliding +1 and sequential nucleosomes away from the TSS and into properly spaced arrays^26^, we believe this evidence supports the conclusion that these promoters with increased pause region Pol II density upon NURF knockdown also have misphased +1 nucleosomes upon NURF knockdown. Therefore, NURF has a role in proper pausing and chromatin architecture in the early gene body, and without the activity of NURF, pause release and the transition to productive elongation are dysregulated (Fig. 1f).

**Figure 3:**
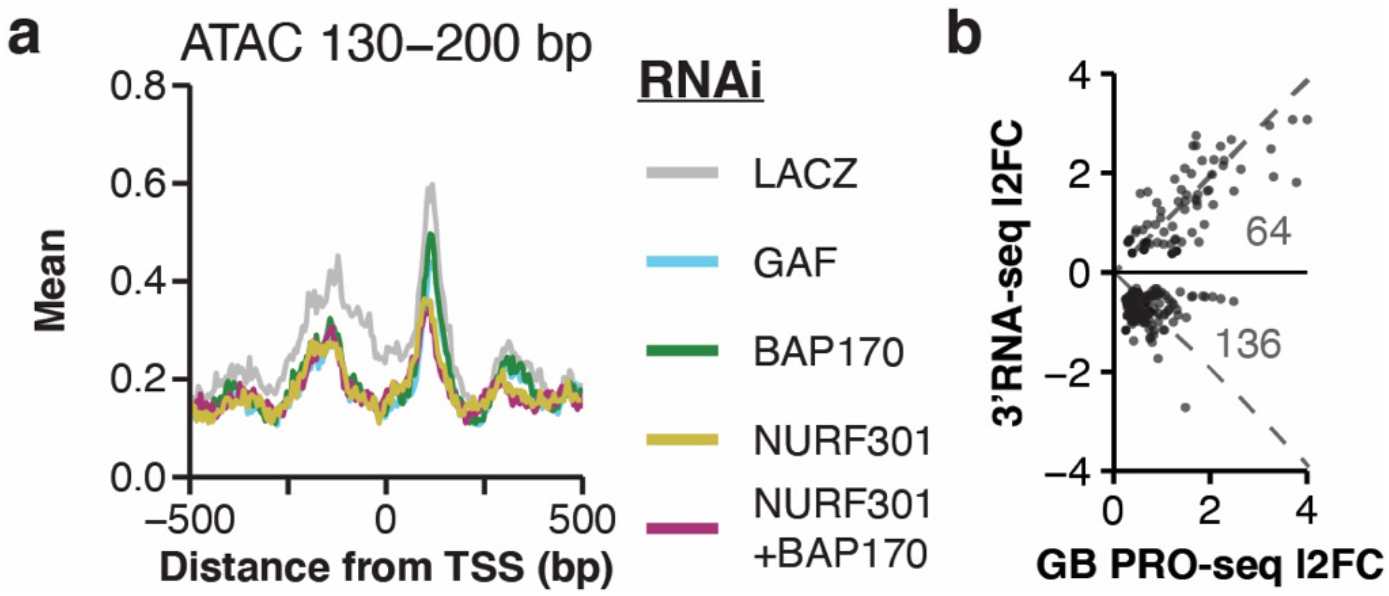
NURF positions nucleosomes which influences pause release and elongation. **(a)** Centers of mononucleosome sized ATAC-seq (fragments 130–200 bp) signal in 1 bp bins at all promoters with NURF301-RNAi upregulated pausing (n=831; see Extended Data Figure 5C; DESeq2 FDR < 0.01). Signal is the mean of 1,000 sub-samplings of 10% of regions. **(b)** PRO-seq gene body (TSS+200–TES-200) log2 fold change compared to RNA-seq log2 fold change (in the last 1 kb region) upon NURF301 RNAi treatment. Only genes with DESeq2 FDR < 0.1 for both PRO-seq and RNA-seq are shown. Number of points in each quadrant are also shown. Dashed lines are 1-to-1 l2FC.

Our model that GAF recruits and functionally synergizes with NURF to ensure efficient pause release and transition to productive elongation predicts that mRNA output would be decreased upon NURF depletion. Indeed, this is observed quite broadly (Extended Data Fig. 5k). GAF interacts physically with NURF^5,19,20^, and GAF-dependent promoters have increased gene body Pol II density by PRO-seq (Fig. 1f, right panel); as such we reasoned that genes with increased gene body Pol II density upon NURF knockdown (n=831) might represent primary targets of NURF. Genes with increased GB PRO-seq signal in the NURF knockdown split into two classes with the majority having decreased mRNA-seq signal (Fig. 3b). This can be explained by Pol II moving more slowly without the activity of NURF, which leads to decreased mRNA output despite increased Pol II density (PRO-seq). Further analysis revealed that genes which increased GB PRO-seq and decreased 3’mRNA-seq upon NURF knockdown (Fig. 3b, bottom half), when compared to those that have increased GB PRO-seq and increased mRNA-seq signal (Fig. 3b, upper half), are normally: (i) less paused; (ii) more expressed; (iii) characterized by higher promoter ATAC-seq hypersensitivity signal that narrows upstream of the TSS upon NURF knockdown; (iv) marked by a well-positioned +1 nucleosome that shows decreased signal upon NURF knockdown; and (v) distinguished by greater gene body polymerase density that further increases upon NURF knockdown (Extended Data Fig. 9b–f, respectively). Taken together, we propose that these findings indicate that these moderately expressed, less paused genes depend more strongly upon the activity of NURF to ensure proper nucleosome positioning. Upon NURF depletion, nucleosomes present an energy barrier to productive elongation, which leads to higher gene body polymerase density despite lower mRNA output as a result of slow-moving polymerases.

We speculate that without the activity of NURF, nucleosomes might drift into sequence-determined “energy wells”—tracts of DNA sequence where nucleosome eviction is less energetically favored—that are difficult for Pol II to transit, especially in the early stage of pause release. Under this model, without the assistance of NURF, both pause release and productive elongation would be inefficient due to the increased energy barrier more tightly DNA-associated nucleosomes present to transcribing Pol II. It was previously demonstrated that in the absence of NURF, +1 nucleosomes drift toward the TSS, and early gene body nucleosomes are mis-phased out to ~1kb at NURF-bound promoters using MNase-seq in *Drosophila* embryonic tissue^26^.

NURF mutant animals have less intense MNase-seq signal associated with the +1 nucleosome at NURF-bound promoters, and the signal maxima shifts ~12 bp towards the TSS^26^. Without NURF, these nucleosomes likely are free to drift into positions that are energetically opposed to Pol II transit, leading to inefficient pause release and therefore increased pause region PRO-seq signal. Taken together, these results indicate that GAF recruits NURF to promoters where it ensures proper nucleosome positioning in the early gene body for energetically favorable nucleosome transit by Pol II, a process downstream of PBAP’s GAF-directed eviction of nucleosomes from NFRs.

## Summary

We demonstrate that GAF is a sequence-specific pioneer factor in *Drosophila*, that depends on the activity of both SWI/SNF (PBAP) and ISWI (NURF) family ATP-dependent nucleosome remodeling complexes to establish optimal chromatin architecture for transcription at target promoters (Fig. 4a). SWI/SNF (PBAP) evicts nucleosomes from promoters, establishing a nucleosome-free region which allows Pol II to be recruited and initiate transcription (Fig. 4b). This first major step of transcription allows Pol II to begin transcription and progress to the promoter-proximal pause region. ISWI (NURF) then ensures that the nucleosomes along the early gene body are properly phased, thereby facilitating Pol II to transition to pause release and productive elongation in an energetically favorable manner (Fig. 4c). This work solidifies decades of *in vitro* biochemistry findings in *Drosophila* by resolving the roles of these factors *in vivo*, and to our knowledge is the first report of a pioneer factor working cooperatively with both ISWI and SWI/SNF remodelers to establish transcription-permissive chromatin at target promoters in metazoans.

**Figure 4:**
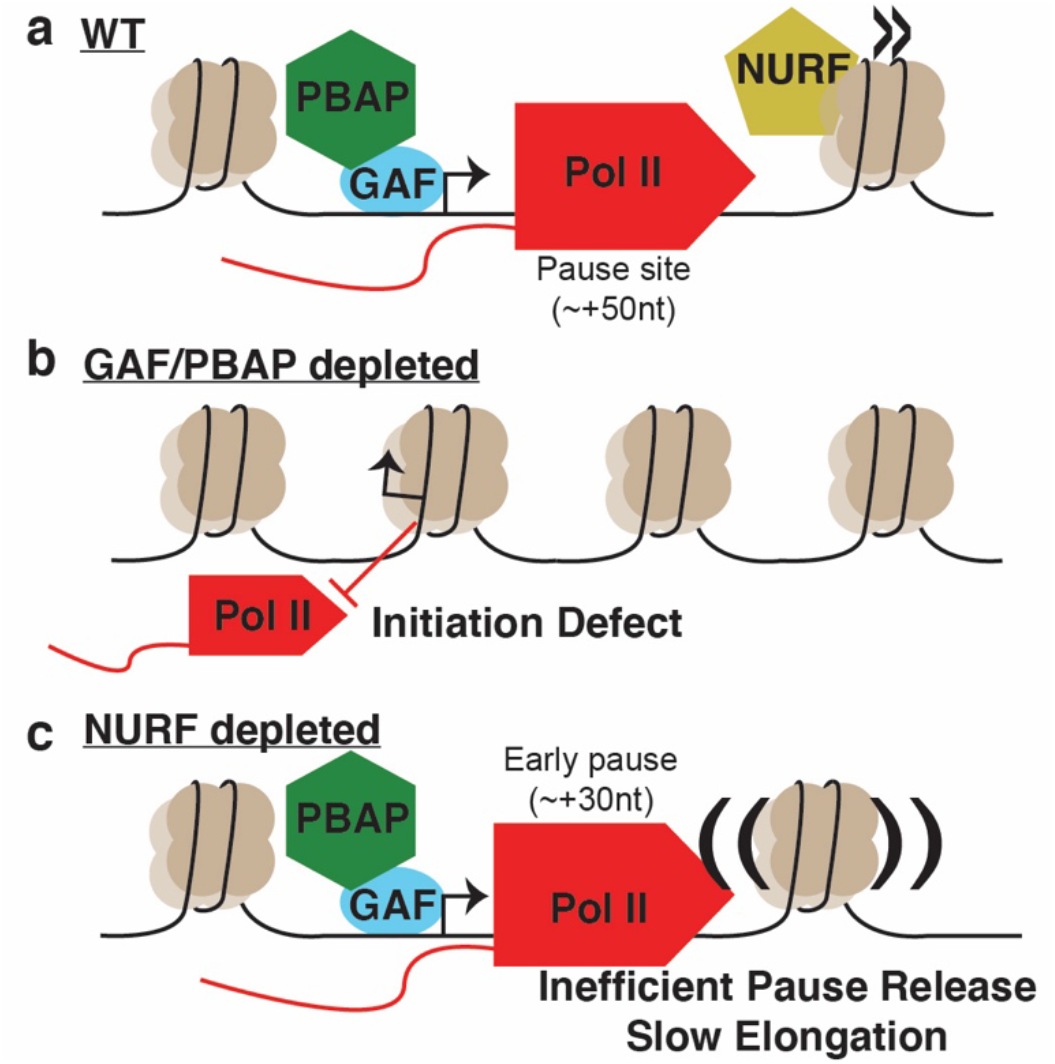
Nucleosome remodelers and pioneer factors coordinate to establish permissive chromatin architecture. **(a–c)** Cartoon summarizing the findings of this article.

These results indicate that a single pioneer transcription factor (GAGA-factor) is able to orchestrate the activity of multiple nucleosome remodeling complexes that regulate the first three stages of the transcription cycle (recruitment, pausing, and transition to productive elongation). GAF and PBAP knockdowns have virtually identical effects on promoter chromatin accessibility and Pol II pausing, showing that GAF synergizes with PBAP to clear promoters of nucleosomes and allow Pol II to be recruited, where it rapidly initiates transcription and traverses to the pause site. However, further analysis demonstrated that GAF also recruits NURF to position the +1 nucleosome, which allows for efficient pause release and transition to productive elongation. The effects of the NURF knockdown are masked by the effects of the GAF/PBAP knockdowns, because without Pol II recruitment, nucleosome positioning along the gene body appears to be mostly irrelevant.

Strikingly, these roles for pioneer factors and specific nucleosome remodeler family members seem to also be consistent with limited recent data in mammals^11,13^, which indicate that this finding might represent a deeply conserved mechanism throughout all of Eukarya. To our knowledge, this is the first report of a single transcription factor with these expansive capabilities in metazoans, and the first view of not only how sequence specific pioneer factors and nucleosome remodelers unite to regulate chromatin, but also how the resulting chromatin structure effects nascent transcription and mRNA production.

## Data availability

All sequencing data has been deposited in GEO (GSE149339). All DESeq2 results tables, raw signal and normalized bigWig files, gene lists, and ATAC-seq peaks can be accessed at https://www.github.com/jaj256/GAF. For ease of viewing, we have also created a custom UCSC track hub with pooled normalized data that can be imported to the UCSC genome browser using this link: https://github.com/JAJ256/GAF/raw/master/hub.txt.

## Code availability

All code used to analyze data and create figures is available at: https://www.github.com/jaj256/GAF.

## Acknowledgements

We would like to thank members of the Lis lab and Feschotte labs for critical reading of the manuscript and helpful discussions regarding data analysis. We thank Paul Badenhorst for open discussion and data-sharing regarding his work on NURF in *Drosophila* embryos. We thank the Cornell BRC Genomics facility and Peter Schweitzer for assistance with Illumina sequencing. We are grateful to Abdullah Ozer for assistance with PRO-cap TSS correction. This work was supported by NIH grants GM025232 and HG009393 to J.T.L. J.J. was supported by NHGRI fellowship F31HG010820. The content is solely the responsibility of the authors and does not necessarily represent the official views of the National Institutes of Health.

## Author contributions

J.J., F.M.D., and J.T.L. conceptualized the study and designed the experimental plan. J.J. performed all experiments and data analysis and wrote the first draft of the manuscript. J.J., F.M.D., and J.T.L. revised the manuscript.

## Methods

### Cell culture and RNAi treatments

*Drosophila* S2 cells were maintained at 25 °C in M3 + BPYE medium with 10% FBS. Two biological replicates were performed for each RNAi treatment as previously described^17^, except dsRNA complementary to LACZ, GAF, BAP170, NURF301, or both BAP170 and NURF301 was added to cultures. We generated dsRNA by PCR amplifying a dsDNA template from S2 genomic DNA with T7 RNA Polymerase promoters on the 5’ end of both strands, and then generated dsRNA using lab-made T7 RNA Polymerase. See Supplementary Table 1 for oligonucleotide primer sequences. All RNAi treatments were done using 10 μg/mL dsRNA, including the BAP170+NURF301 condition (5 μg/mL each). After 5 days, an equal volume of 25 °C serum-free M3 + BYPE was added to cultures and they were incubated at 25 °C for 20 min (this was to mimic a paired heat-stress experiment that was performed alongside these experiments but is not presented in this publication). Cells were then harvested for PRO-seq, ATAC-seq, and 3’RNA-seq, and aliquots were lysed by boiling in 1x Laemmli buffer for western blot analysis.

### Western blots

Western blots were performed using anti-GAF (lab-made, 1:500) or anti-NURF301(Novus Biologicals Cat. No. 40360002; 1:100), with anti-Chromator (lab-made, 1:2000) as a loading control. Loading was standardized by cell number and for each RNAi treatment, a serial 2-fold dilution curve was analyzed compared to the LACZ-RNAi condition. Protein was detected using dual-color secondary antibodies and blots were imaged using the LI-COR Odyssey system.

### Custom genomes

To facilitate accurate counting of spike-in reads, published PRO-seq data that did not contain spiked-in human cells^17^, was aligned to a repeat-masked human genome (hg38 assembly^27^, retrieved from the UCSC genome browser^28^) using bowtie2^29^ using default parameters. Unique alignments (mapq > 1) were retained, and any regions with alignments were masked using bedtools maskfasta^30^. This custom-masked genome was then combined with the *Drosophila* genome (dm6 genome assembly^31^, retrieved from the UCSC genome browser^28^. This allowed us to align PRO-seq data (containing both human- and fly-derived sequences) to this combined genome and ensured that no drosophila-derived reads aberrantly mapped to the human genome and skewed spike-in normalization factors. We also masked any region in the dm6 genome assembly larger than 100 bp with greater than 80% homology to *Hsp70Aa* in order to uniquely map sequencing data to a single copy of *Hsp70*.

### Gene annotations

We started with a list of all unique FlyBase transcripts^32^, and reassigned the TSS based on the site of maximum PRO-cap signal^33^ in the window of TSS ± 50 bp. We then filtered out transcripts less than 500 bp long and removed any duplicate transcripts (occasionally two isoforms with TSSs within 50 bp of each other are corrected to the same PRO-cap maximum site, resulting in a duplicate transcript). We then discarded any transcript for which length-normalized PRO-seq signal in the TSS-upstream region (−400 to −100) was more than half that in the pause region (−50 to +100) or more than that in the gene body region (TSS+200 to TES-200). This removed transcripts for which read-through transcription from an upstream gene is a major driver of signal within that gene and removes most transcript isoforms other than the most expressed isoform. This filtering left a list of 9,375 genes, which was the starting point for DESeq2^34^ differential expression testing and PCA analyses.

### PRO-seq library preparation

PRO-seq library preparation was performed as previously described^21,33^ using 2 × 10^7^ cells per condition. We spiked in 2.7 × 10^5^ human K562 cells immediately after harvesting cells to facilitate robust normalization of PRO-seq data. We substituted MyOne C1 Streptavidin beads for the M280 beads recommended by the published protocol, as their negatively charged surface is thought to reduce non-specific nucleic acid binding, and we used 5’ and 3’ adapters that each had a 6N unique molecular identifier at the ligation junction to facilitate computational PCR deduplication of reads. PRO-seq libraries were all amplified for 11 PCR cycles and sequenced on an Illumina NextSeq in 37×37 paired end mode.

### PRO-seq data analysis

Data quality was assessed with fastqc^35^. Adapters were trimmed and UMIs were extracted using fastp^36^, and rRNA reads were removed using bowtie2^29^. Reads were then aligned to the combined dm6/hg38 genome assembly described above and reads aligning uniquely (mapq > 10) to the human genome were counted for spike-in normalization. Reads were then mapped to the dm6 genome using bowtie2^29^, and only uniquely mapping reads (mapq > 10) were retained. Alignments were PCR-deduplicated using UMI-tools^37^ (spike-in alignments were also de-duplicated). BigWig coverage tracks of alignment 3’ end positions in single base pair bins were then generated using deepTools^38^. Normalization factors were derived by taking the minimum number of reads mapped to the spike-in genome across all samples and dividing that by the number of mapped spike-in reads for each sample (Supplementary Equation 1). The alignment pipeline used can be found at http://github.com/jaj256/PROseq_alignment.sh, commit 55a08db). See Supplementary Table 2 for PRO-seq alignment metrics and normalization factors.

### ATAC-seq library preparation

ATAC-seq was performed as previously described^39^, with some modifications for *Drosophila* cells. Briefly, 10^5^ cells were washed with ice-cold PBS, and then resuspended in ice-cold lysis buffer (10 mM Tris-Cl pH 7.4, 10 mM NaCl, 3 mM MgCl_2_, 0.1% NP-40, and 1x Pierce Protease Inhibitors [Thermo Scientific]) and incubated on ice for 3 min. Nuclei were then pelleted and resuspended in Transposition buffer (10 mM Tris-Cl pH 7.4, 10% DMF, and 5 mM MgCl_2_), and 1.5 μL of lab-made Tn5 transposase was added. After a 30 min incubation in a thermomixer at 37 °C, DNA was extracted using phenol:chloroform, PCR amplified for 11 cycles, and sequenced on an Illumina NextSeq in 37×37 paired end mode.

### ATAC-seq data analysis

Reads were aligned to the dm6 genome assembly using bowtie2^29^ in local mode, and only unique alignments were retained (mapq > 10). Signal was then divided into two classes: hypersensitivity (paired end alignments with fragment size < 120 bp, which represents hypersensitive chromatin and generates fragments smaller than mononucleosomes), and mononucleosome (paired end alignments with fragment size 130–200 bp, which represents two transposition events that roughly flank a mononucleosome sized region). Coverage tracks were generated using deepTools^38^. For hypersensitivity signal, entire alignments were “piled up” to generate coverage tracks, and for mononucleosome data only the central 3 bp of each alignment were considered. ATAC-seq peaks were called using macs2^40^. See Supplementary Table 3 for alignment metrics.

### 3’RNA-seq

3’RNA-seq libraries were prepared using the QuantSeq 3’ mRNA-seq Library Prep Kit (Lexogen) with the UMI add-on kit. For each condition, 10^6^ cells were added to a fixed amount of ERCC Spike-In RNA Mix (Invitrogen), and RNA was extracted using TRIzol reagent (Invitrogen). RNA treated with RNase free DNase I (Thermo Scientific), and the absence of DNA was confirmed using the Qubit dsDNA-HS assay (Thermo Scientific). RNA quality was confirmed using denaturing agarose gel electrophoresis. 3’RNA-seq libraries were prepared using 325 ng of total RNA per condition according to manufacturer instructions and sequenced on an Illumina NextSeq in 75 bp single end mode. Reads were trimmed of adapter and polyA sequences and UMIs were extracted using fastp^36^. Reads were then aligned to a combined dm6/ERCC reference genome using STAR^41^, and reads mapped to the ERCC standards were counted for spike-in normalization. Alignments were PCR-deduplicated using UMI-tools^37^, and only unique reads were retained (mapq = 255). The 5’ ends of reads were used to generate signal tracks (so that transcripts were scored in a read-length independent manner) using deepTools^38^. Spike-in normalization factors were calculated as described above for PRO-seq. See Supplemental Table 4 for alignment metrics and normalization factors.

### CUT&RUN

CUT&RUN was performed as described^42,43^. We used both anti-GAF (lab-made) or anti-NURF301(Novus Biologicals Cat. No. 40360002) at a 1:10 dilution for the antibody binding step. ProteinA-MNase was incubated with calcium on ice for 30 minutes, and cleaved fragments were recovered by phenol:chloroform extraction. Library prep was performed using the following steps: (i) Ends of digested fragments were repaired by incubation for 30 min at 25 °C with 0.5 U/μL T4 PNK, 0.12 U/μL T4 DNA Polymerase, and 0.05 U/μL Klenow Fragment in 1X T4 DNA ligase buffer (with ATP) and 0.5 mM dNTPs; (ii) Fragments were A-tailed by incubation for 30 min at 37 °C with 0.25 U/μL Klenow exo- and 0.5 mM dATP in 1X NEBuffer 2; (iii) Adapters were added by incubation on the lab bench for 2 h with 4.38 nM lab-made Illumina TruSeq forked adapters and 24 U/μL T4 DNA ligase in 1X T4 DNA Ligase buffer (with ATP); (iv) library DNA was recovered using AMPure XP beads (1.8X concentration) and PCR amplified for 15 cycles (all enzymes from New England Biolabs). Libraries were sequenced on an Illumina NextSeq in 37×37 paired end mode. Adapters sequences were removed using fastp^36^, and reads were aligned to the dm6 reference genome using bowtie2^29^. Only uniquely mapped reads (mapq > 10) with fragment size smaller than 120 bp were retained, and signal coverage tracks were generated using deepTools^38^. Signal was normalized per million mapped reads. See Supplementary Table 5 for alignment metrics.

### Reanalysis of Published Data

GAF ChIP-seq^16^ raw reads were downloaded and mapped to the dm6 genome assembly using bowtie2^29^, and only uniquely mapping reads were retained (mapq > 10). Single end reads were extended 200 bp and reads-per-million normalized coverage tracks were generated using deepTools^38^. Peaks were called using macs2^40^. A M1BP ChIP-seq^23^ signal track was downloaded and converted for the dm6 assembly using liftOver^28^, and signal was normalized on a per-million basis. M1BP-knockdown PRO-seq^17^ normalized signal tracks were accessed and converted to the dm6 genome assembly as above. BEAF-32 ChIP-seq^25^ raw reads were downloaded, aligned using bowtie2^29^, and only uniquely mapping reads were retained (mapq > 10). Single end reads were extended 200 bp and reads-per-million normalized coverage tracks were generated using deepTools^38^. See Supplementary Table 6 for accession numbers for all published data used in this manuscript.

### DE testing

Signal counting in each set of regions for each data type was performed using functions from the BRGenomics package^44^. Differential expression testing and principal component analysis was performed using DESeq2^34^. Genes with adjusted p-value < 0.01 were considered differentially expressed. See Supplemental Code 6 for details.

### Browser shots

Browser shots were generated using a custom R function, which can be found at https://github.com/JAJ256/browser_plot.R (commit 1352d5c).

### Metaprofiles

Metaprofiles were generated using the BRGenomics package^44^ by calculating a signal matrix across all regions in a set using the bin size specified, then sampling 10% of regions 1000 times to calculate the mean and 75% confidence interval. In some cases, confidence intervals were removed to avoid over-plotting. Visualization was performed using ggplot2^45^.

### Motif analysis

To search for motifs overrepresented in one set of promoters compared to another, we used DREME^46^ with an e-value threshold of 0.001.

### Classification of GAF-bound promoters

We considered a promoter GAF-bound if the promoter region (−500 to TSS) overlapped with a GAF ChIP-seq peak (see above). We then considered these GAF-bound promoters as having GAF-dependent pausing or GAF-independent pausing on the basis of whether or not they had significantly decreased PRO-seq in the pause region compared to the LACZ-RNAi control (DESeq2 FDR < 0.01, log2 Fold Change < 0). We further subdivided the GAF-bound promoters with GAF-independent pausing by whether they were bound by M1BP, BEAF-32, or both, with “bound” defined as falling within the top 25% of promoters in our total set of GAF-bound promoters with GAF-independent pausing when rank-ordered by total ChIP-seq signal within the promoter (−500 to TSS) for a given factor. Heatmaps were created using the ComplexHeatmap R package^47^.

## Extended Data Figures

**Extended Data Figure 1:**
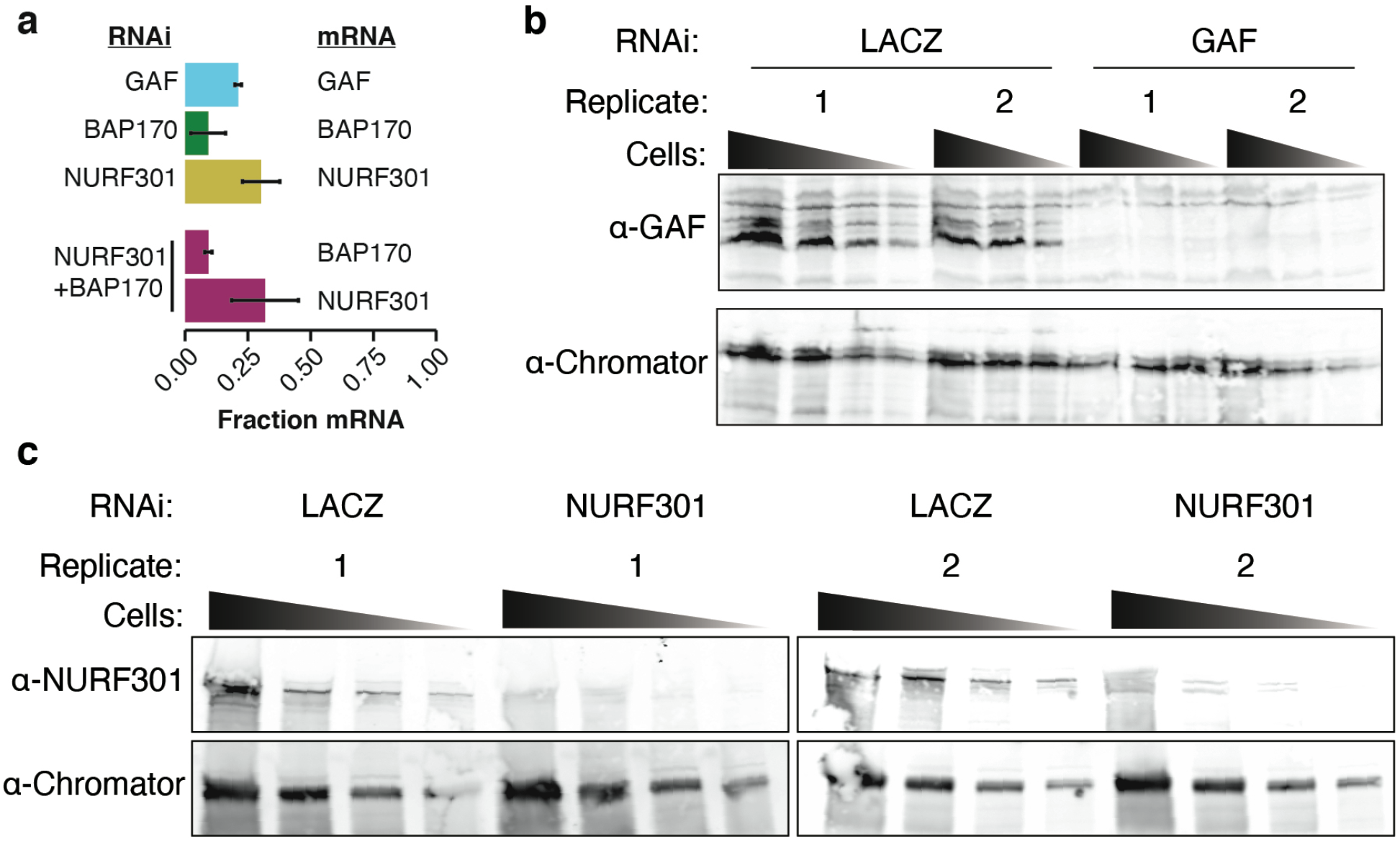
Knockdown efficiency of GAF-RNAi and NURF301-RNAi by western blot. **(a)** Fraction mRNA remaining compared to the LACZ control for each knockdown target. *BAP170* and *NURF301* transcript levels are displayed independently for the double knockdown. Bar height is mean; error bars are SD. **(b)** Western blot showing the amount of GAF protein remaining after 5 days of RNAi treatment compared to the LACZ-RNAi control. For each condition a serial 2-fold dilution series of cells was loaded. Chromator is included as a loading control. **(c)** Western blot showing the amount of NURF301 protein remaining after 5 days of RNAi treatment compared to the LACZ-RNAi control. For each condition a serial 2-fold dilution series of cells was loaded. Chromator is included as a loading control.

**Extended Data Figure 2:**
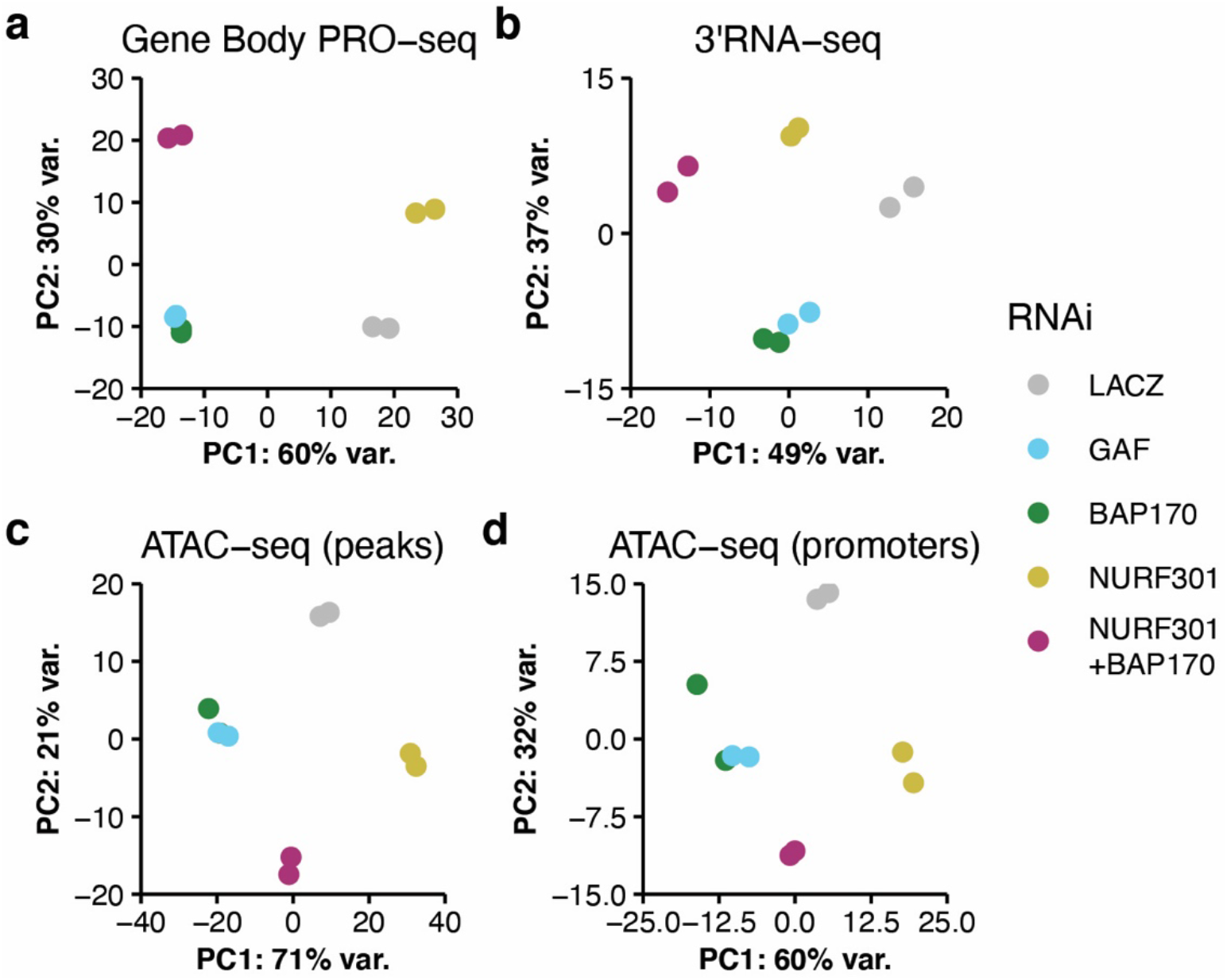
Principal component analysis of gene body PRO-seq, 3’RNA-seq, and ATAC-seq. **(a)** Principal component analysis of spike-in normalized PRO-seq signal in the gene body (TSS+200 to TES-200) of genes in a filtered list (see methods, n=9,375). **(b)** Principal component analysis of spike-in normalized 3’RNA-seq signal of genes in a filtered list (n=9,375), signal was counted in the last 1 kb region of each gene. **(c)** Principal component analysis of library size-normalized ATAC-seq signal in ATAC-seq peak summits ± 100 bp (n=39,806). **(d)** Principal component analysis of library size-normalized ATAC-seq signal in promoter regions (−1000–TSS) of genes in a filtered list (n=9,375).

**Extended Data Figure 3:**
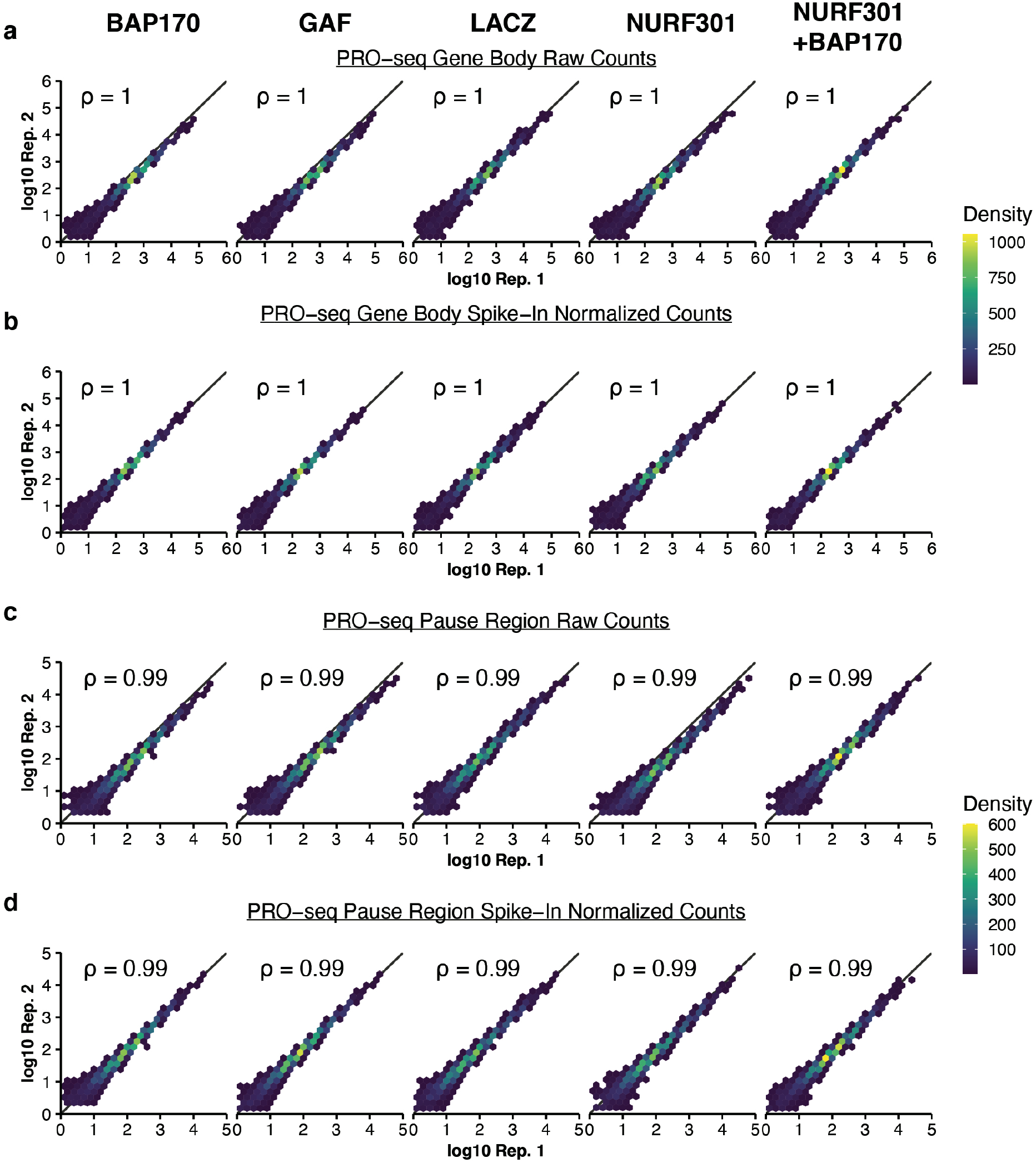
Correlation and normalization of PRO-seq data. **(a)** Raw number of PRO-seq reads mapped to the gene body (TSS+200 to TES-200) of each gene in a filtered list (n=9,375, see methods) for all conditions, with replicate 1 on the x-axis and replicate 2 on the y-axis. ρ is Spearman’s rho. To accommodate overplotting, the plot space was divided into hexbins and color-mapped by the number of genes in each bin. **(b)** As in **(a)**, but counts were normalized using spike-in scaling factors. **(c)** As in **(a)**, but raw number of PRO-seq reads mapped to the pause region (−50 to +100) of each gene. **(d)** As in **(c)**, but counts were normalized using spike-in scaling factors.

**Extended Data Figure 4:**
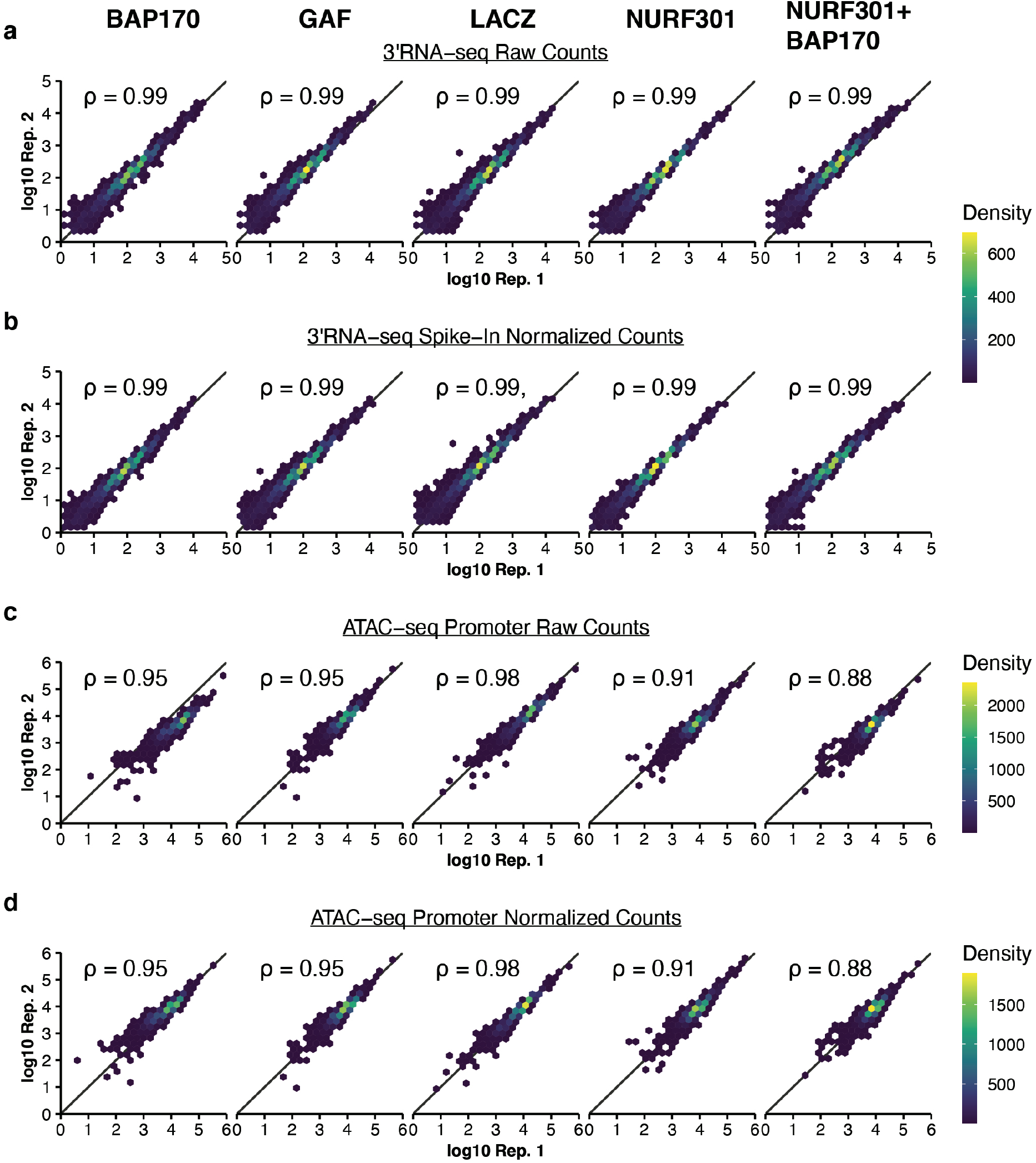
Correlation and normalization of RNA-seq and ATAC-seq data. **(a)** Raw number of 3’RNA-seq reads mapped to the last 1 kb region of each gene in a filtered list (n=9,375, see methods) for all conditions, with replicate 1 on the x-axis and replicate 2 on the y-axis. ρ is Spearman’s rho. To accommodate overplotting, the plot space was divided into hexbins and color-mapped by the number of genes in each bin. **(b)** As in **(a)**, but counts were normalized using spike-in scaling factors. **(c)** As in **(a)**, but raw number of ATAC-seq reads (< 120bp) mapped to the promoter region (−1000– TSS) of each gene. **(d)** As in **(c)**, but counts were normalized using library-size scaling factors (DESeq2).

**Extended Data Figure 5:**
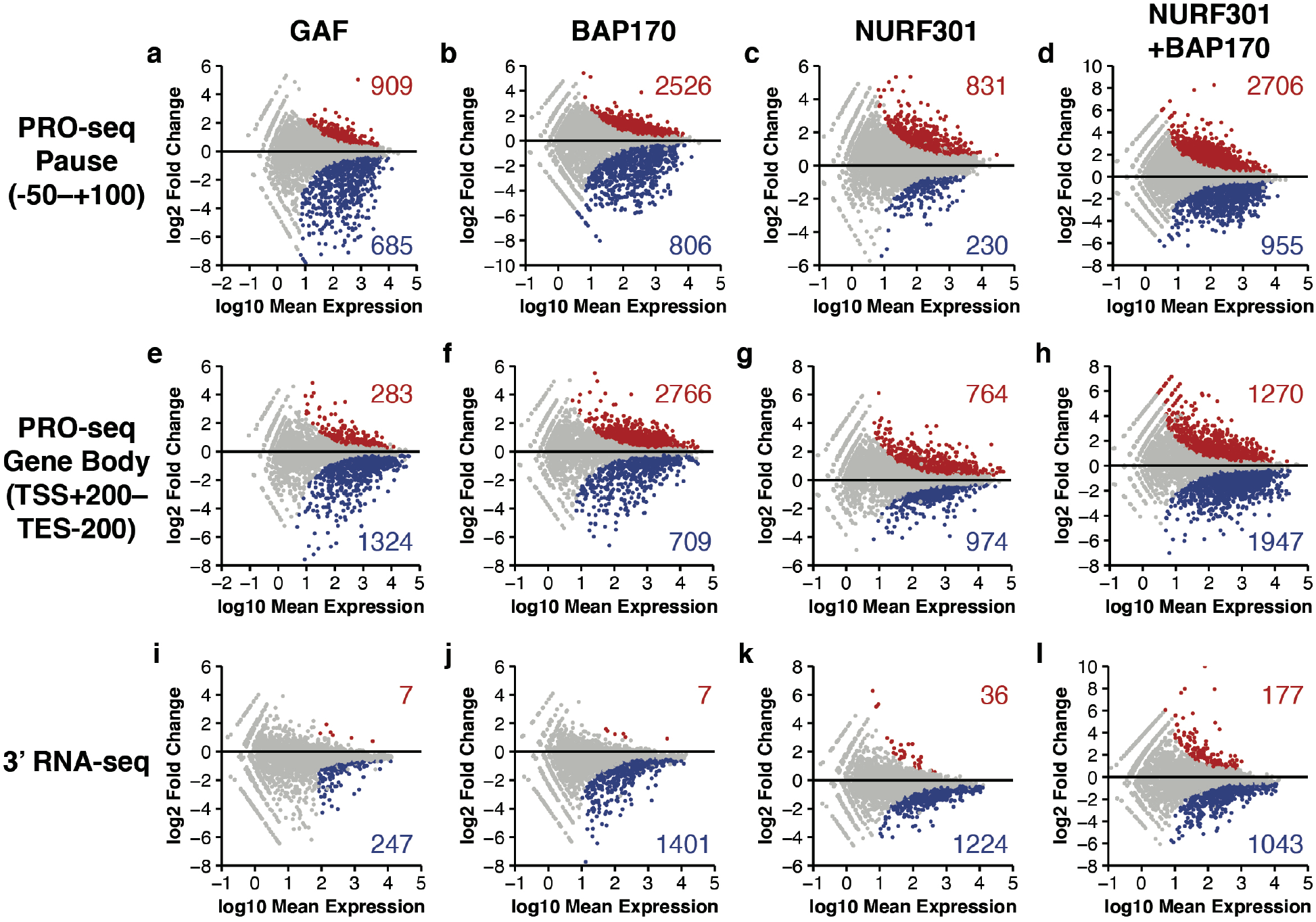
MA plots for all conditions and data types. **(a)** MA plot for the comparison of spike-in normalized GAF-RNAi vs. LACZ-RNAi PRO-seq in the pause region (TSS −50 to +100); DESeq2 FDR < 0.01. Number of genes significantly up- or down-regulated is also shown. **(b)** As in **(a)** but BAP170 RNAi vs. LACZ-RNAi. **(c)** As in **(a)** but NURF301 RNAi vs. LACZ-RNAi. **(d)** As in **(a)** but NURF301+BAP170 RNAi vs. LACZ-RNAi. **(e)** MA plot for the comparison of spike-in normalized GAF-RNAi vs. LACZ-RNAi PRO-seq in the gene body (TSS+200–TES-200); DESeq2 FDR < 0.01. Number of genes significantly up- or down-regulated is also shown. **(f)** As in **(e)** but BAP170 RNAi vs. LACZ-RNAi. **(g)** As in **(e)** but NURF301 RNAi vs. LACZ-RNAi. **(h)** As in **(e)** but NURF301+BAP170 RNAi vs. LACZ-RNAi. **(i)** MA plot for the comparison of spike-in normalized GAF-RNAi vs. LACZ-RNAi 3’RNA-seq signal in the last 1 kb region of each gene; DESeq2 FDR < 0.01. Number of genes significantly up- or down-regulated is also shown. **(j)** As in **(i)** but BAP170 RNAi vs. LACZ-RNAi. **(k)** As in **(i)** but NURF301 RNAi vs. LACZ-RNAi. **(l)** As in **(i)** but NURF301+BAP170 RNAi vs. LACZ-RNAi.

**Extended Data Figure 6:**
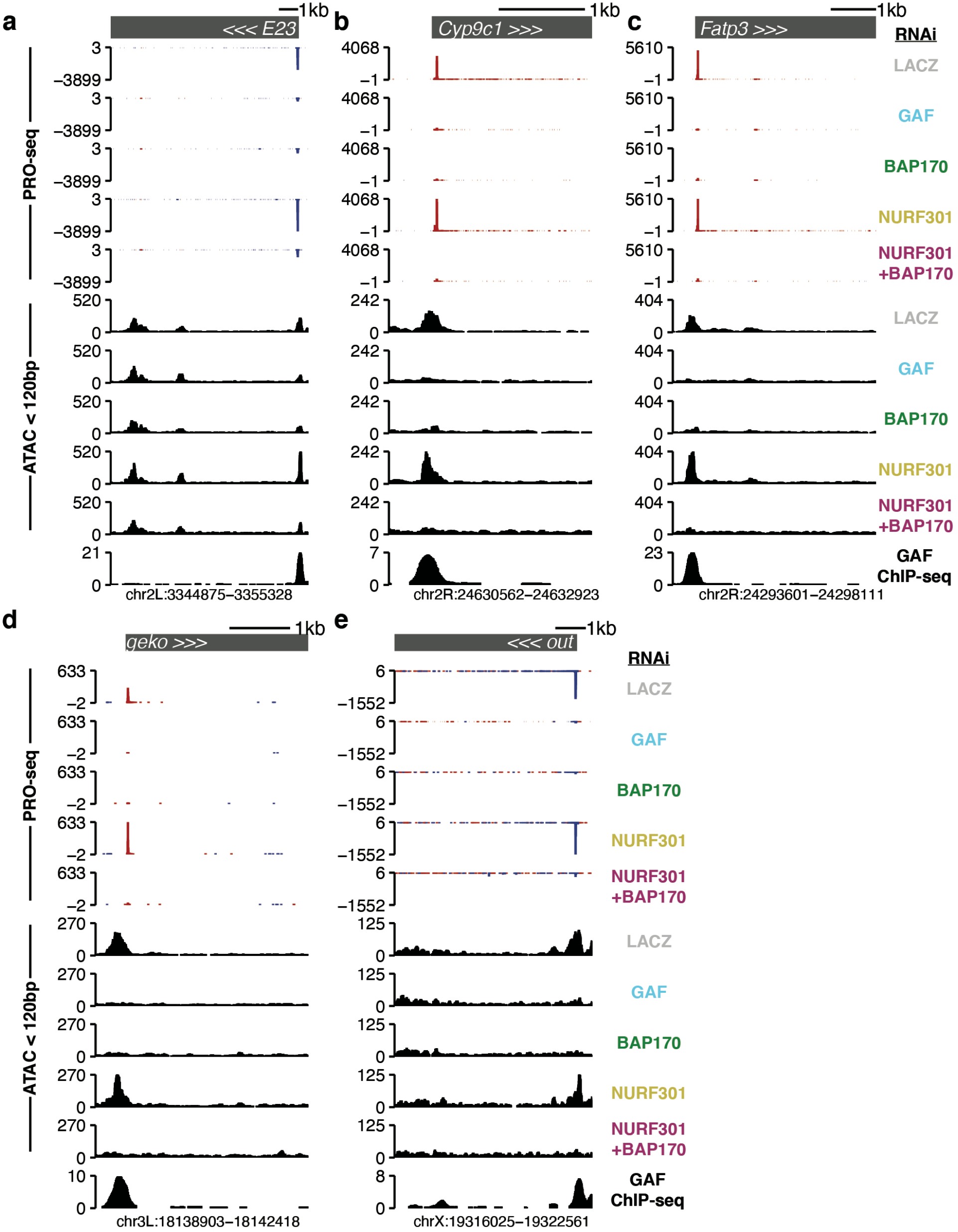
Examples of GAF-dependent promoters. **(a)** Browser shot of a gene with GAF-dependent pausing (*E23-RC*). Tracks are each shown as the mean of two replicates. ATAC-seq was filtered to retain only paired-end alignments with insert size < 120 bp. **(b)** As in **(a)** but *Cyp9c1-RA*. **(c)** As in **(a)** but *Fatp3-RA*. **(d)** As in **(a)** but *geko-RB*. **(e)** As in **(a)** but *out-RA*.

**Extended Data Figure 7:**
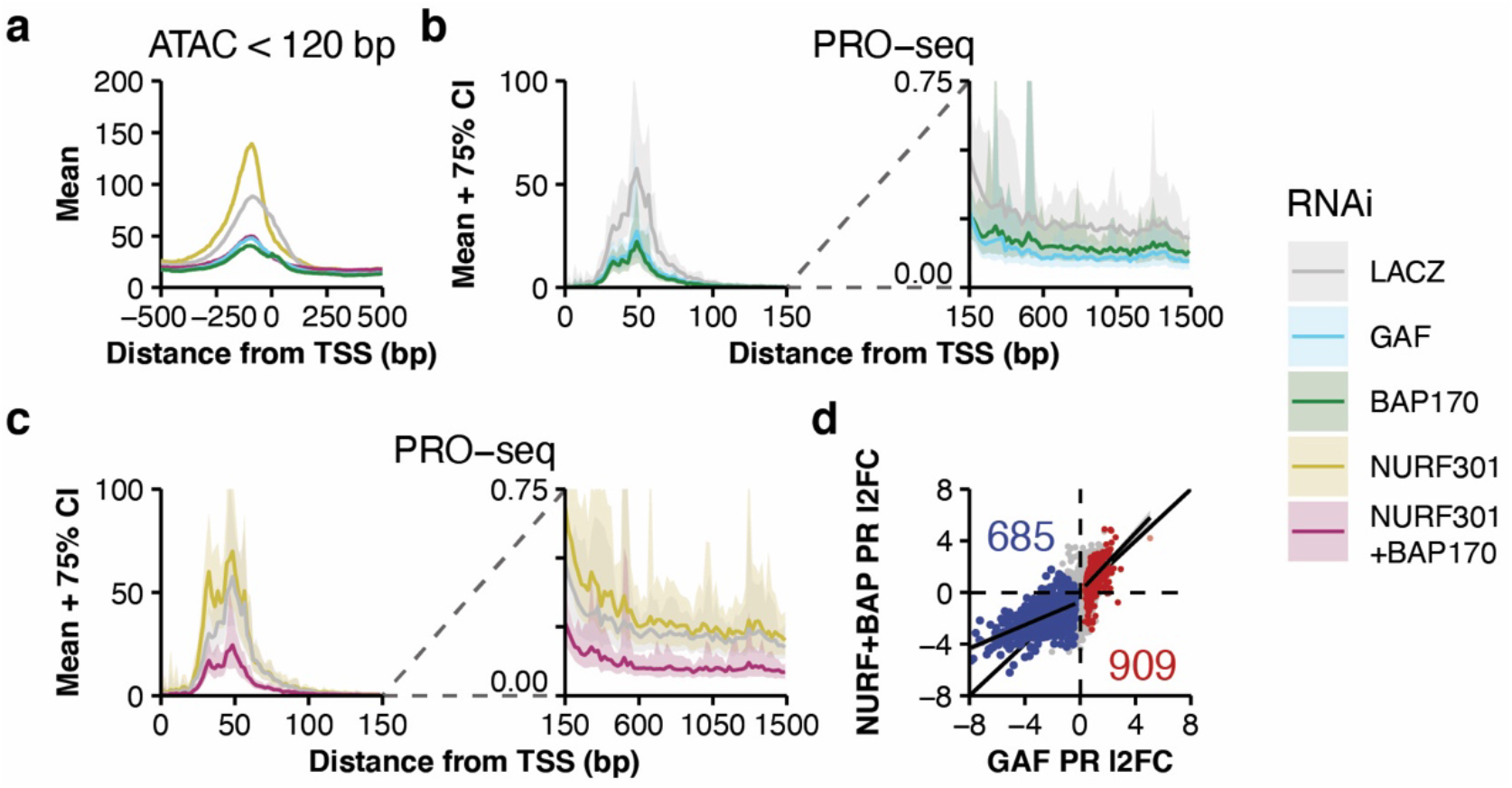
PBAP opens promoter chromatin in coordination with GAF. **(a)** ATAC-seq (< 120 bp) signal for each RNAi treatment in 1 bp bins across all promoters with significantly decreased PRO-seq signal in the pause region (−50–+100) upon BAP170-RNAi treatment (n=806; see Extended Data Figure 5b; DESeq2 FDR < 0.01). Signal is the mean of 1,000 sub-samplings of 10% of regions. **(b)** PRO-seq signal at genes with BAP170-dependent pausing (n=806) for the LACZ, GAF, and BAP170 RNAi treatments. The pause region (left panel) is in 2 bp bins, and the first 1.5 kb of the gene body (right panel) is in 20 bp bins. Data is shown as mean (solid line) and 75% confidence interval (shaded area), derived from 1,000 sub-samplings of 10% of regions. **(c)** As in **(b)**, but for LACZ, NURF301, and NURF301+BAP170 RNAi treatments. **(d)** Pause region (TSS −50 to +100) PRO-seq log2 fold change after GAF-RNAi treatment compared to NURF301+BAP170-RNAi. Red and blue points are significantly up- or down-regulated upon GAF-RNAi treatment (DESeq2 FDR < 0.01). The line and shaded area are a GLM and 95% confidence interval fit to significantly up- or down-regulated genes.

**Extended Data Figure 8:**
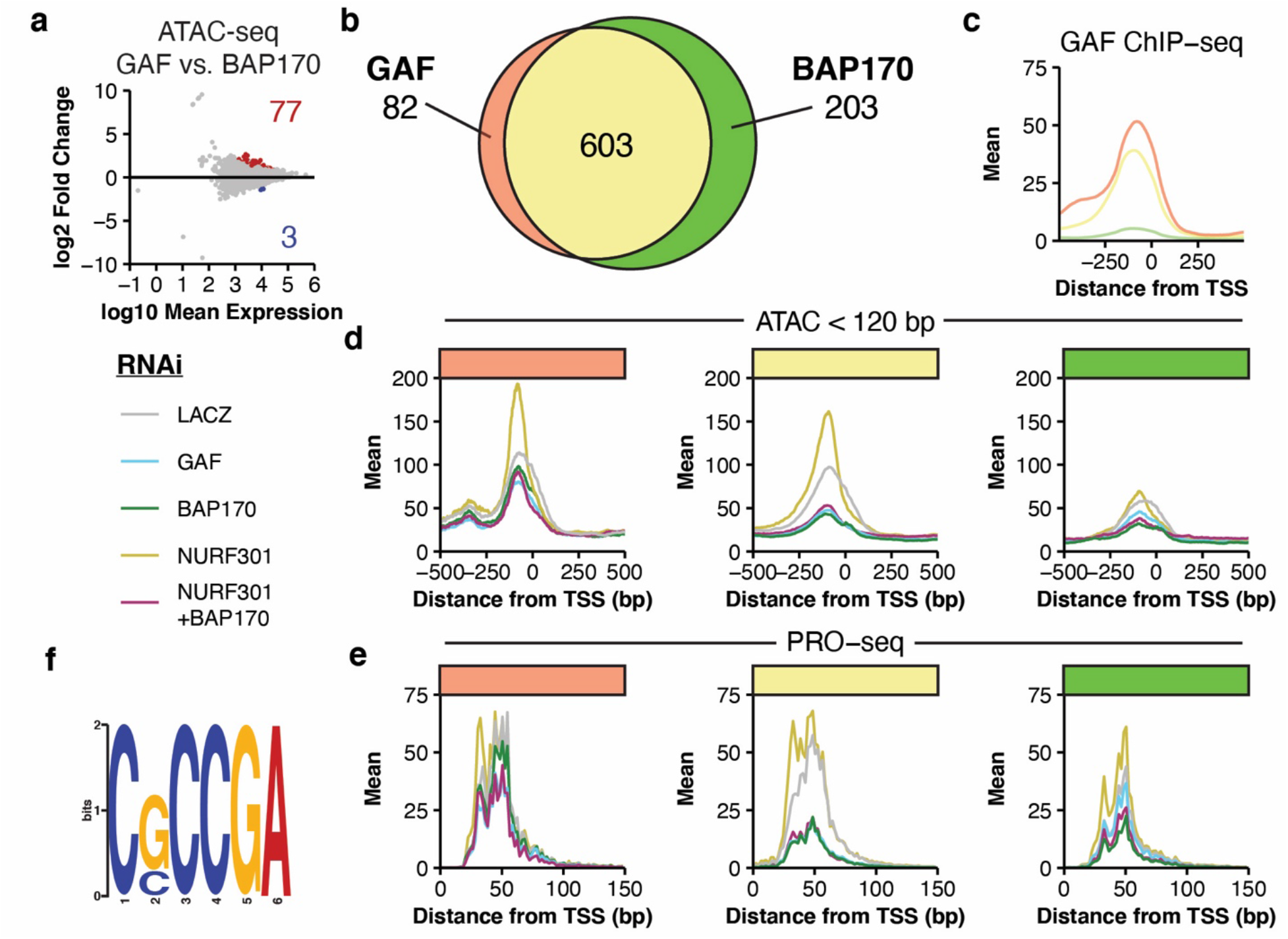
Effects of GAF and BAP-170 RNAi are indistinguishable. **(a)** MA plot for the comparison of GAF-RNAi vs. BAP170-RNAi ATAC-seq (fragments < 120 bp) in ATAC-seq peak summits ± 100 bp (n=39,806); DESeq2 FDR < 0.01. Number of regions significantly up- or down-regulated is also shown. **(b)** Intersection of promoters significantly downregulated by either GAF-RNAi (n=685) or BAP170-RNAi (n=806). **(c)** Promoter GAF ChIP-seq signal in 5 bp bins at gene sets color coded as in **(b)**. **(d)** Promoter ATAC-seq (fragments < 120 bp) signal for each RNAi treatment in 1 bp bins at gene sets defined in **(b)**. Signal is the mean of 1,000 sub-samplings of 10% of regions. **(e)** Pause region PRO-seq signal for each RNAi treatment in 2 bp bins at gene sets defined in **(b)**. Signal is the mean of 1,000 sub-samplings of 10% of regions. **(f)** Motif enriched in BAP170 exclusive promoters (−500 to TSS; n=203; green in **(b)**–**(e)**) over GAF/BAP170 dependent promoters (−500 to TSS; n=603; yellow in **(b)**–**(e)**). DREME E-value < 0.001.

**Extended Data Figure 9:**
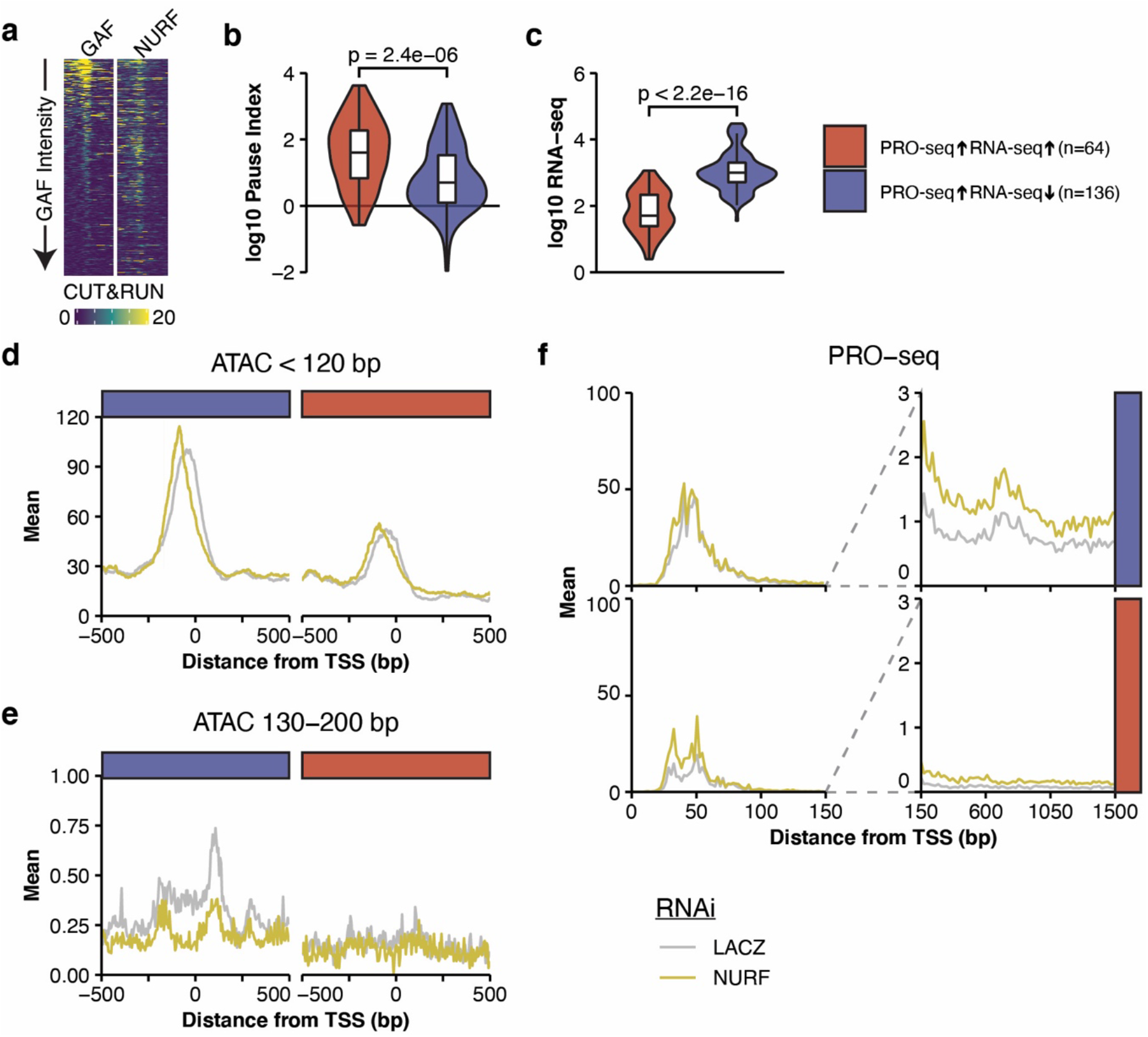
NURF-RNAi causes bifurcated changes in nascent transcription and mRNA levels. **(a)** GAF and NURF301 CUT&RUN signal at the top quartile of genes by GAF CUT&RUN signal (TSS ± 500). Rows are sorted by GAF signal and sort order is maintained. **(b)** Distribution of Pause Indices (pause region PRO-seq signal / length normalized gene body PRO-seq signal) for genes with increased gene body PRO-seq density and increased RNA-seq signal (DESeq FDR < 0.1; n=64) or genes with increased gene body PRO-seq density but decreased RNA-seq signal (n=136). See Figure 3a for gene classes. **(c)** Distribution of RNA-seq normalized counts for the two classes of genes described in **(b)**. **(d)** ATAC-seq (fragments < 120 bp) signal for NURF301 and LACZ RNAi treatments in 1 bp bins across promoters of the genes described in **(b)**. Signal is the mean of 1,000 subsamplings of 10% of regions. **(e)** As in **(e)**, but ATAC-seq fragments 130–200 bp. Only the central 3 bp of each alignment were considered when generating signal tracks. **(f)** PRO-seq signal for NURF301 and LACZ RNAi treatments at genes described in **(b)**. The pause region (left panel) is in 2 bp bins, and the first 1.5 kb of the gene body (right panel) is in 20 bp bins. Signal is the mean of 1,000 sub-samplings of 10% of regions. In **(b)** and **(c)** filled violins represent the distribution and boxplots show the median (center line), 25% and 75% quartiles (hinges), and 1.5*IQR (whiskers). Outliers are not plotted, and p-value is from a Mann-Whitney U test.

## References

1. Zaret, K. S. & Mango, S. E. Pioneer transcription factors, chromatin dynamics, and cell fate control. Curr. Opin. Genet. Dev. 37, 76–81 (2016).

2. Mayran, A. & Drouin, J. Pioneer transcription factors shape the epigenetic landscape. J. Biol. Chem. 293, 13795–13804 (2018).

3. Vallot, A. & Tachibana, K. The emergence of genome architecture and zygotic genome activation. Curr. Opin. Cell Biol. 64, 50–57 (2020).

4. Moshe, A. & Kaplan, T. Genome-wide search for Zelda-like chromatin signatures identifies GAF as a pioneer factor in early fly development. Epigenetics Chromatin 10, 33 (2017).

5. Xiao, H. et al. Dual functions of largest NURF subunit NURF301 in nucleosome sliding and transcription factor interactions. Mol. Cell 8, 531–543 (2001).

6. Tsukiyama, T. & Wu, C. Purification and properties of an ATP-dependent nucleosome remodeling factor. Cell 83, 1011–1020 (1995).

7. Vihervaara, A., Duarte, F. M. & Lis, J. T. Molecular mechanisms driving transcriptional stress responses. Nat. Rev. Genet. 19, 385–397 (2018).

8. Kubik, S., Bruzzone, M. J. & Shore, D. Establishing nucleosome architecture and stability at promoters: Roles of pioneer transcription factors and the RSC chromatin remodeler. Bioessays 39, (2017).

9. Krietenstein, N. et al. Genomic Nucleosome Organization Reconstituted with Pure Proteins. Cell 167, 709–721.E12 (2016).

10. Wagner, F. R. et al. Structure of SWI/SNF chromatin remodeller RSC bound to a nucleosome. Nature 579, 448–451 (2020).

11. Hainer, S. J., Bošković, A., McCannell, K. N., Rando, O. J. & Fazzio, T. G. Profiling of Pluripotency Factors in Single Cells and Early Embryos. Cell 177, 1319–1329.E11 (2019).

12. He, S. et al. Structure of nucleosome-bound human BAF complex. Science 367, 875–881 (2020).

13. King, H. W. & Klose, R. J. The pioneer factor OCT4 requires the chromatin remodeller BRG1 to support gene regulatory element function in mouse embryonic stem cells. Elife 6, (2017).

14. Farkas, G. et al. The Trithorax-like gene encodes the Drosophila GAGA factor. Nature 371, 806–808 (1994).

15. Wilkins, R. C. & Lis, J. T. GAGA factor binding to DNA via a single trinucleotide sequence element. Nucleic Acids Res. 26, 2672–2678 (1998).

16. Fuda, N. J. et al. GAGA factor maintains nucleosome-free regions and has a role in RNA polymerase II recruitment to promoters. PLoS Genet. 11, e1005108 (2015).

17. Duarte, F. M. et al. Transcription factors GAF and HSF act at distinct regulatory steps to modulate stress-induced gene activation. Genes Dev. 30, 1731–1746 (2016).

18. Tsukiyama, T. & Wu, C. Purification and properties of an ATP-dependent nucleosome remodeling factor. Cell 83, 1011–1020 (1995).

19. Lomaev, D. et al. The GAGA factor regulatory network: Identification of GAGA factor associated proteins. PLoS One 12, e0173602 (2017).

20. Nakayama, T., Shimojima, T. & Hirose, S. The PBAP remodeling complex is required for histone H3.3 replacement at chromatin boundaries and for boundary functions. Development 139, 4582–4590 (2012).

21. Mahat, D. B. et al. Base-pair-resolution genome-wide mapping of active RNA polymerases using precision nuclear run-on (PRO-seq). Nat. Protoc. 11, 1455–1476 (2016).

22. Gilchrist, D. A. et al. Pausing of RNA polymerase II disrupts DNA-specified nucleosome organization to enable precise gene regulation. Cell 143, 540–551 (2010).

23. Li, J. & Gilmour, D. S. Distinct mechanisms of transcriptional pausing orchestrated by GAGA factor and M1BP, a novel transcription factor. EMBO J. 32, 1829–1841 (2013).

24. Yang, J., Ramos, E. & Corces, V. G. The BEAF-32 insulator coordinates genome organization and function during the evolution of Drosophila species. Genome Res. 22, 2199–2207 (2012).

25. Liang, J. et al. Chromatin immunoprecipitation indirect peaks highlight long-range interactions of insulator proteins and Pol II pausing. Mol. Cell 53, 672–681 (2014).

26. Kwon, S. Y., Grisan, V., Jang, B., Herbert, J. & Badenhorst, P. Genome-Wide Mapping Targets of the Metazoan Chromatin Remodeling Factor NURF Reveals Nucleosome Remodeling at Enhancers, Core Promoters and Gene Insulators. PLoS Genet. 12, e1005969 (2016).

## References (Methods)

27. Lander, E. S. et al. Initial sequencing and analysis of the human genome. Nature 409, 860–921 (2001).

28. Haeussler, M. et al. The UCSC Genome Browser database: 2019 update. Nucleic Acids Res. 47, D853–D858 (2019).

29. Langmead, B. & Salzberg, S. L. Fast gapped-read alignment with Bowtie 2. Nat. Methods 9, 357–359 (2012).

30. Quinlan, A. R. & Hall, I. M. BEDTools: A flexible suite of utilities for comparing genomic features. Bioinformatics 26, 841–842 (2010).

31. Hoskins, R. A. et al. The Release 6 reference sequence of the Drosophila melanogaster genome. Genome Res. 25, 445–458 (2015).

32. Thurmond, J. et al. FlyBase 2.0: the next generation. Nucleic Acids Res. 47, D759–D765 (2019).

33. Kwak, H., Fuda, N. J., Core, L. J. & Lis, J. T. Precise maps of RNA polymerase reveal how promoters direct initiation and pausing. Science. 339, 950–953 (2013).

34. Love, M. I., Huber, W. & Anders, S. Moderated estimation of fold change and dispersion for RNA-seq data with DESeq2. Genome Biol. 15, 550 (2014).

35. Andrews, S. FastQC: A quality control tool for high throughput sequence data. (2010). Available at: http://www.bioinformatics.babraham.ac.uk/projects/fastqc/.

36. Chen, S., Zhou, Y., Chen, Y. & Gu, J. fastp: an ultra-fast all-in-one FASTQ preprocessor. Bioinformatics 34, i884–i890 (2018).

37. Smith, T., Heger, A. & Sudbery, I. UMI-tools: modeling sequencing errors in Unique Molecular Identifiers to improve quantification accuracy. Genome Res. 27, 491–499 (2017).

38. Ramírez, F., Dündar, F., Diehl, S., Grüning, B. A. & Manke, T. deepTools: a flexible platform for exploring deep-sequencing data. Nucleic Acids Res. 42, W187–W191 (2014).

39. Buenrostro, J. D., Wu, B., Chang, H. Y. & Greenleaf, W. J. ATAC-seq: A Method for Assaying Chromatin Accessibility Genome-Wide. Curr. Protoc. Mol. Biol. 109, 21.29.1–21.29.9 (2015).

40. Zhang, Y. et al. Model-based analysis of ChIP-Seq (MACS). Genome Biol. 9, (2008).

41. Dobin, A. et al. STAR: ultrafast universal RNA-seq aligner. Bioinformatics 29, 15–21 (2013).

42. Skene, P. J. & Henikoff, S. An efficient targeted nuclease strategy for high-resolution mapping of DNA binding sites. Elife 6, (2017).

43. Skene, P. J., Henikoff, J. G. & Henikoff, S. Targeted in situ genome-wide profiling with high efficiency for low cell numbers. Nat. Protoc. 13, 1006–1019 (2018).

44. DeBerardine, M. BRGenomics: Tools for the efficient analysis of high-resolution genomics data. R package version 0.99.31. (2020). doi:10.18129/B9.bioc.BRGenomics

45. Wickham, H. ggplot2: Elegant Graphics for Data Analysis. (Springer-Verlag, New York, 2016).

46. Bailey, T. L. DREME: Motif discovery in transcription factor ChIP-seq data. Bioinformatics 27, 1653–1659 (2011).

47. Gu, Z., Eils, R. & Schlesner, M. Complex heatmaps reveal patterns and correlations in multidimensional genomic data. Bioinformatics 32, 2847–2849 (2016).

