## Supplemental Tables 1-6 for "Pioneer factor GAF cooperates with PBAP and NURF to regulate transcription"

### Supplemental Information for 'Pioneer factor GAF cooperates with PBAP and NURF to regulate transcription'

### Table of Contents

1. Supplementary Table 1. Oligonucleotides used for generating RNAi templates
2. Supplementary Table 2. PRO-seq alignment metrics
3. Supplementary Table 3. ATAC-seq alignment metrics
4. Supplementary Table 4. 3'RNA-seq alignment metrics
5. Supplementary Table 5. CUT&RUN alignment metrics
6. Supplementary Table 6. External data sources
7. Supplementary Equation 1. Spike-In normalization strategy

All computer code used to perform analyses in this paper is available here: https://github.com/JAJ256/GAF

#### Supplementary Table 1. Oligonucleotides used for generating RNAi templates

| Name | Sequence |
|---|---|
| LACZ-fwd | GAATTAATACGACTCACTATAGGGAGAGATATCCTGCTGATGAAGC |
| LACZ-rev | GAATTAATACGACTCACTATAGGGAGAGCAGGAGCTCGTTATCGC |
| GAF-fwd | GAATTAATACGACTCACTATAGGGATGGTTATGTTGGCTGGCGTCAA |
| GAF-rev | GAATTAATACGACTCACTATAGGGATCTTTACGCGTGGTTTGCGT |
| BAP170-fwd | GAATTAATACGACTCACTATAGGGTGGACGGAATAGAGCTACCTGG |
| BAP170-rev | GAATTAATACGACTCACTATAGGGTCAATGGAGCGAGAGGTGG |
| NURF301-fwd | GAATTAATACGACTCACTATAGGGATGTTGAATACTGGTTGACATAGTTC |
| NURF301-rev | GAATTAATACGACTCACTATAGGGAGTGCTAATCCGGCATGATA |

#### Supplementary Table 2. PRO-seq alignment metrics

| RNAi | Rep. | Raw Reads | % Adapter | % rRNA | % PCR Dups | Uniq. Non-Dup | Scale Factor |
|---|---|---|---|---|---|---|---|
| BAP170 | 1 | 39,244,087 | 6.63% | 22.05% | 0.01% | 15,379,401 | 0.626245 |
| BAP170 | 2 | 26,110,434 | 8.35% | 20.05% | 0.01% | 9,304,431 | 1.000000 |
| GAF | 1 | 48,692,340 | 6.43% | 19.10% | 0.02% | 20,159,679 | 0.364060 |
| GAF | 2 | 29,022,776 | 5.40% | 20.27% | 0.01% | 11,329,566 | 0.650773 |
| LACZ | 1 | 40,319,830 | 7.44% | 22.28% | 0.01% | 16,451,927 | 0.533495 |
| LACZ | 2 | 35,740,698 | 5.82% | 20.88% | 0.01% | 14,594,540 | 0.579465 |
| NURF301+BAP170 | 1 | 45,591,479 | 8.56% | 16.97% | 0.01% | 14,763,333 | 0.522120 |
| NURF301+BAP170 | 2 | 38,224,454 | 7.09% | 16.19% | 0.01% | 12,152,986 | 0.556803 |
| NURF301 | 1 | 61,102,403 | 7.90% | 10.55% | 0.01% | 17,854,770 | 0.559650 |
| NURF301 | 2 | 34,094,767 | 7.09% | 10.40% | 0.01% | 9,103,431 | 0.964054 |

#### Supplementary Table 3. ATAC-seq alignment metrics

| RNAi | Rep. | Raw Reads | Uniq. Aligned Reads |
|---|---|---|---|
| BAP170 | 1 | 38,500,034 | 30,231,402 |
| BAP170 | 2 | 14,483,253 | 11,502,911 |
| GAF | 1 | 21,962,543 | 17,108,608 |
| GAF | 2 | 22,832,804 | 17,912,219 |
| LACZ | 1 | 23,978,897 | 19,095,473 |
| LACZ | 2 | 18,977,303 | 15,141,294 |
| NURF301+BAP170 | 1 | 24,629,540 | 19,568,654 |
| NURF301+BAP170 | 2 | 17,109,780 | 13,816,496 |
| NURF301 | 1 | 22,298,270 | 17,243,075 |
| NURF301 | 2 | 18,414,735 | 14,401,208 |

#### Supplementary Table 4. 3'RNA-seq alignment metrics

| RNAi | Rep. | Raw Reads | % Adapter | % PCR Dups | Uniq. Non-Dup | ERCC Reads | Scale Factor |
|---|---|---|---|---|---|---|---|
| BAP170 | 1 | 9,897,071 | 4.75% | 68.53% | 3,114,663 | 113,089 | 0.696478 |
| BAP170 | 2 | 12,083,788 | 4.17% | 71.77% | 3,411,719 | 122,112 | 0.645014 |
| GAF | 1 | 10,572,796 | 5.76% | 71.71% | 2,991,223 | 104,509 | 0.753658 |
| GAF | 2 | 16,673,438 | 4.67% | 74.68% | 4,222,261 | 131,954 | 0.596905 |
| LACZ | 1 | 14,553,663 | 5.35% | 72.75% | 3,966,094 | 105,724 | 0.744996 |
| LACZ | 2 | 13,133,097 | 9.67% | 73.64% | 3,461,387 | 78,764 | 1.000000 |
| NURF301+BAP170 | 1 | 12,153,411 | 5.52% | 74.60% | 3,086,672 | 111,297 | 0.707692 |
| NURF301+BAP170 | 2 | 22,904,765 | 9.37% | 76.98% | 5,271,654 | 160,525 | 0.490665 |
| NURF301 | 1 | 12,878,694 | 5.98% | 73.75% | 3,380,145 | 117,876 | 0.668194 |
| NURF301 | 2 | 10,521,245 | 5.22% | 70.39% | 3,115,527 | 101,947 | 0.772598 |

#### Supplementary Table 5. CUT&RUN alignment metrics

| Target | Raw Reads | % Adapter | Uniq. Aligned Reads |
|---|---|---|---|
| GAF | 18,499,020 | 2.34% | 12,097,380 |
| NURF301 | 17,234,208 | 2.13% | 10,713,798 |

#### Supplementary Table 6. External data sources

| Description | PMID | GEO | SRA | Remapped? |
|---|---|---|---|---|
| M1BP ChIP-seq | 23708796 | GSE49842 | SRP028808 | No |
| BEAF-32 ChIP-seq | 24486021 | GSE52962 | SRP033490 | Yes |
| GAF ChIP-seq | 25815464 | GSE40646 | SRP015432 | Yes |
| M1BP PRO-seq | 27492368 | GSE77607 | SRP069335 | No |

#### Supplementary Equation 1. Spike-In normalization strategy

$$Signal[i]_{normalized} = Signal[i]_{raw} \cdot \frac{min\{SpikeIn_i \; ... \; SpikeIn_n\}}{SpikeIn_i}$$

Where:

$Signal[i]_{normalized}$ = Normalized signal for sample $i$

$Signal[i]_{raw}$ = Raw counts for sample $i$

$min\{SpikeIn_i \; ... \; SpikeIn_n\}$ = Minimum number spike-in reads mapped across all samples

$SpikeIn_i$ = Number of spike-in reads mapped for sample $i$
